# Tracking polyhydroxyalkanoate biosynthesis in thermophilic microorganisms

**DOI:** 10.1101/2025.05.06.652502

**Authors:** Brendan Schroyen, Radwa Moanis, Hannelore Geeraert, Niko Van den Brande, Ulrich Hennecke, Stanislav Obruča, Iva Buchtíková, Karel Sedlář, Petr Sedláček, Eveline Peeters

## Abstract

Polyhydroxyalkanoates are biopolyesters synthesized and stored in intracellular granules by diverse prokaryotes. Despite intense research efforts and prior evidence of a rather widespread phylogenetic occurrence of the related genetic machinery, reports on extreme thermophilic and hyperthermophilic polyhydroxyalkanoates producers remain scarce. However, thermophilic cell factories for bioplastic production would serve as an excellent example of Next-Generation Industrial Biotechnology. In this study, we aim to address this research gap by establishing a bioinformatics pipeline to mine genomes of extremely and moderately thermophilic microorganisms for signatures of potential polyhydroxyalkanoate production. Based on a collection of verified protein sequences of polyhydroxyalkanoate polymerase PhaC, the key biosynthetic enzyme, carefully curated sets of thermophilic bacterial and archaeal genomes were screened. This revealed that although PhaC-encoding genes are prevalent in diverse moderately thermophilic bacteria, they are absent in the considered extreme thermophilic bacteria. In contrast, a few limited examples of extreme thermophilic archaea were found to encode putative *phaC* genes embedded within a typical polyhydroxyalkanoate synthesis operon in their genomes, namely within the genera *Ferroglobus*, *Geoglobus* and *Archaeoglobus*, while no hits were found in extreme thermophilic bacteria. The latter included *Thermus thermophilus*, which was previously reported as a polyhydroxyalkanoates producer. This was refuted in our bioinformatics analysis and moreover, the predicted absence of polyhydroxyalkanoates synthesis in *T. thermophilus* was experimentally confirmed by employing various extraction and analytical methods. Based on the findings in this study, we conclude that polyhydroxyalkanoate production is very scarce in extreme thermophiles and hyperthermophiles, for reasons that remain to be elucidated.

**Highlights:**

- A bioinformatics pipeline was constructed to screen thermophilic genomes for PhaC.
- PHA production is widespread in moderate thermophiles but rare in extreme thermophiles.
- Extreme thermophilic archaea belonging to specific genera exceptionally harbor PHA synthesis genes.
- No PHA synthesis genes were found in extreme thermophilic bacteria like *Thermus* spp.
- Experimental work confirmed the absence of PHAs in *Thermus thermophilus*.

**Graphical abstract:** 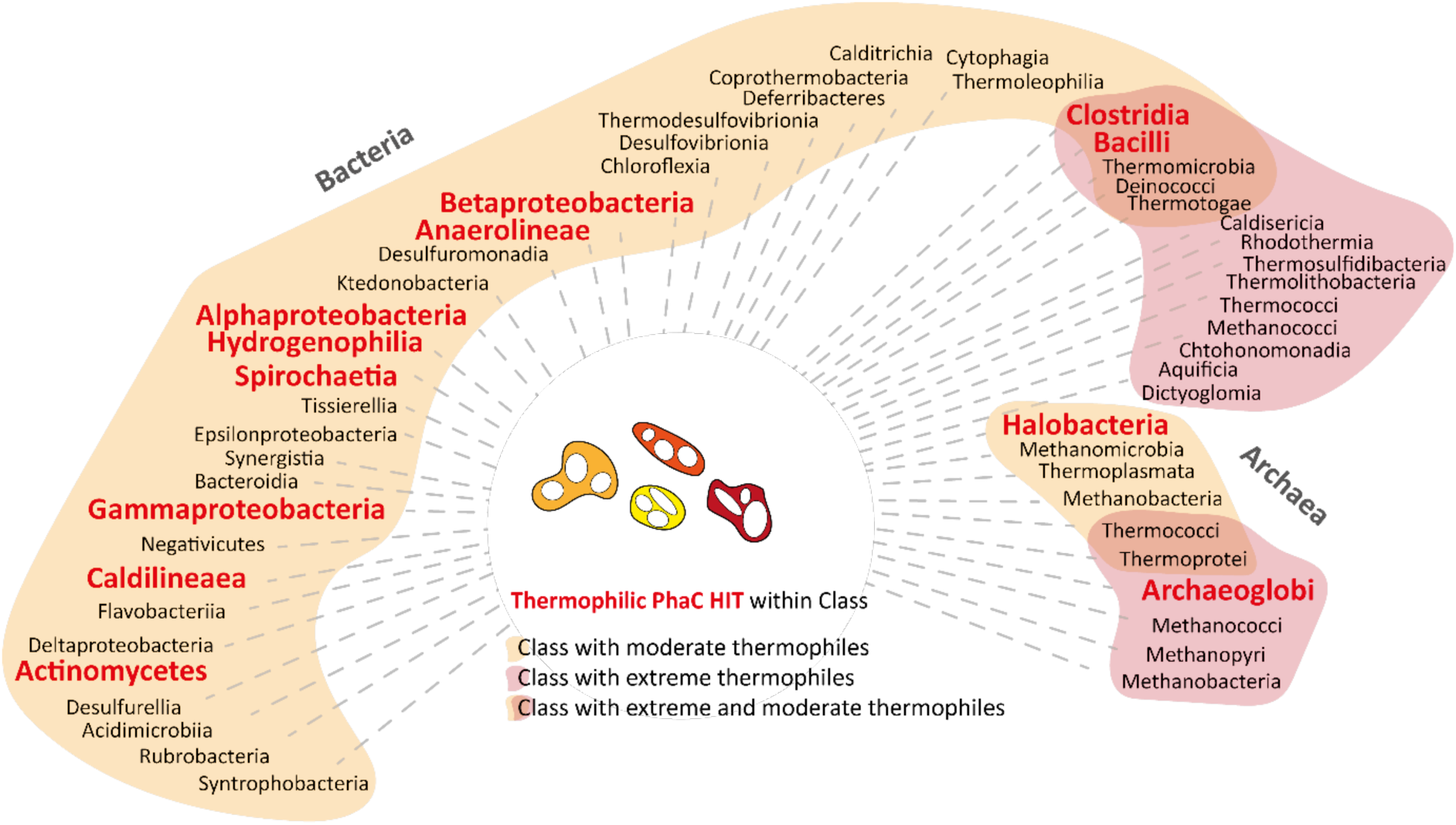

## 1. Introduction

Diverse prokaryotic microbes synthesize polyhydroxyalkanoates (PHAs), polyesters that are stored in intracellular water-insoluble granules [1]. PHA synthesis is intricately connected to the central metabolism and reversible, often taking place simultaneously with degradation, thus leading to a dynamic “PHA cycle” [2]. Initially, PHA synthesis was recognized solely as a carbon and energy reserve strategy of the microbial cell, enabling the capture of organic carbon in conditions of nutritional imbalances thereby supporting growth in environmentally unfavorable conditions [3]. Recent studies point to more versatile biological roles for PHA metabolism, ranging from a strategy to steer central metabolism in developmental processes such as sporulation in *Bacillus* spp. [4] to having complex stress response functions against osmolarity, temperature, pH, UV irradiation and/or oxidative stress conditions [5,6,7]. For example, PHA granules could act as internal scaffolds protecting cells from cellular lysis while the granule-associated proteins could serve as chaperones that protect biomolecules. In addition, the intracellular pool of PHA monomers that are produced might have an extremolyte function, which is especially relevant in conditions of high osmolarity.

Different PHA biosynthetic pathways exist that each involve an array of PHA-specific proteins, besides being connected to central metabolic pathways, such as the β oxidation and fatty acid synthase II pathways [1]. Regardless of species-specific differences in PHA synthesis pathways, all PHA-producing strains harbor a PHA synthase enzyme named PhaC, which catalyzes the polymerization reaction itself [8]. PhaC enzymes can be divided into at least four classes depending on substrate specificity, subunit composition and catalytic activity: while Class I and II PhaC enzymes consist of a single subunit and prefer short chain-length (*scl*) and medium chain-length (*mcl*) PHA monomers, respectively, Class III and IV enzymes are heterodimers consisting of two subunits, with the latter class also catalyzing alcohol cleavage of the PHA polymer chain [9]. Genes contributing to PHA metabolism are often organized in operon-like structures and/or clustered in the genome, although gene synteny is typically not conserved [10].

From a biotechnological perspective, the interest in microbial PHA synthesis is rising, given that the properties of PHA polyesters are comparable with those of petrochemical-based plastics thereby offering a more sustainable and environmentally friendly alternative [1]. Despite this promise, wide industrialization of PHA production has not yet been adopted given that the average production cost is still 4 to 5 times higher than that of petroleum-based plastics [11]. One of the proposed strategies to improve cost-efficiency of microbial PHA production is the use of extremophilic strains, framed in the concept of Next-Generation Industrial Biotechnology, coined by Chen & Jiang [7,8,12]. Indeed, many halophilic and thermophilic species can be cultivated on industrially relevant low-cost feedstocks and enable the establishment of a consolidated process with up- and downstream process steps, such as PHA extraction. Moreover, their use leads to reduced energy costs due to decreased contamination risks, which lowers sterility requirements and reduces the need for energy-intensive cooling [8,13,14].

While the phylogenetic distribution of PHA synthesis in halophilic bacteria and archaea is quite well understood, this is more enigmatic for thermophiles [8]. Indeed, only a limited number of thermophiles (optimal growth temperature (T_opt_) between 45 and 65°C), extreme thermophiles (T_opt_ between 65 and 80°C) and hyperthermophiles (T_opt_ exceeding 80°C) [15,16] have been reported in literature to produce PHAs, with equally limited information on the underlying genetic and metabolic pathways [17]. In early 2000s, several reports stated that the extreme thermophilic bacterial strain *Thermus thermophilus* HB8 synthesizes PHAs [18,19,20]. Recently, a literature study was performed, as well as a bioinformatics analysis of the genomic presence of PhaC-encoding genes, with the aim of broadly mapping bacterial thermophilic PHA producers [8]. This study revealed that most PHA-producing species, either predicted or phenotypically validated, have a T_opt_ in the range between 40 and 60°C, and are thus classified as either thermotolerant or (moderately) thermophilic [8]. Gram-negative thermotolerant or thermophilic PHA-producing species include those that belong to the *Chelatococcus*, *Tepidimonas* and *Caldimonas* genera [8], as well as the thermotolerant *Paracoccus* species *Paracoccus kondratievae* [21]. *Caldimonas thermodepolymerans* is of particular interest because of its catabolic versatility, capable of utilizing xylose and cellobiose, relevant for industrial feedstocks, and because it produces a poly(3-hydroxybutyrate-*co*-3-hydroxyvalerate) co-polymer [22,23,24]. Among Gram-positive bacteria, the *Aneurinibacillus* genus comprises several thermotolerant and thermophilic species that produce PHAs, which have unique capabilities given an exceptional PHA biosynthesis versatility [8,25].

In this study, we aim to close the gap in knowledge with regards to the occurrence of PHA production in extreme thermophilic prokaryotes. A bioinformatics pipeline is developed by using prior knowledge on the highly conserved lipase-like box in PhaC enzymes and by screening a comprehensive set of moderate to extreme and hyperthermophilic bacterial and archaeal genomes. Results were further refined by analyzing other highly conserved PhaC residues, as well as PhaC’s catalytic triad [26]. In addition, we aimed to solve the conundrum of the capability of *T. thermophilus* to synthesize PHAs by performing extractions in different strains, including *T. thermophilus* HB8, followed by chemical characterization. Overall, we demonstrate that PHA synthesis is very scarce in extreme thermophilic and hyperthermophilic genera and speculate on possible reasons that have impeded the evolutionary expansion of PHA metabolism beyond mesophilic and moderate thermophilic species.

## 2. Materials and methods

### 2.1. Bioinformatics screening for putative PhaC homologs in thermophilic genomes

The Comprehensive Enzyme Information System (BRENDA database) [27] and the Kyoto Encyclopedia of Genes and Genomes (KEGG) databases [28] were used to identify relevant pathways in which (putative) enzymes are active. Protein sequences were obtained by browsing the Uniprot Knowledgebase (UniprotKB) database [29] and consulting the Protein database of the National Center of Biotechnology Information (NCBI). Sequences that were manually annotated and part of the Swiss-Prot sublibrary were preferentially selected. In addition, the NCBI Protein database was screened for homologs by using the text query ‘Polyhydroxyalkanoate synthase’ OR ‘PhaC’ [30]. A final list of 16, well-described or automatically annotated, protein sequences covering all four PhaC enzyme classes was assembled (**Supplementary Table S1**).

An annotated list of 137 moderate (17 archaeal and 120 bacterial genera) and 74 extreme thermophiles (36 archaeal and 38 bacterial genera) was manually created based on previously published work [31]. This implies that in total, 11,396 genomes on the species-level were interrogated for the moderate thermophiles (2689 archaeal and 8707 bacterial), and 1,694 genomes for the extreme thermophiles (860 archaeal and 834 bacterial) (**Supplementary File S1**).

Homologs of PhaC proteins were predicted using the online version of Domain Enhanced Lookup Time Accelerated BLAST (DELTA-BLAST) [32] tool available at the NCBI website with the PhaC protein sequence of *Chelatococcus daeguensis* TAD1 (AN: WP_071923939.1) as a query. The DELTA-BLAST analysis was performed separately for each category — moderate thermophilic Bacteria, moderate thermophilic Archaea, extreme thermophilic Bacteria, and extreme thermophilic Archaea — using the genomes belonging to the genera in each group. Standard algorithm parameters were used and the top-5000 hits, of each category, were retained for the next steps.

Multiple sequence alignments were created using the online version of Constraint-based Multiple Alignment Tool (COBALT) [33], hosted on the NCBI website using the standard algorithm parameters. Conservation schemes were created using the ConSurf web server [34]. Finally, putative operons were analysed with operon_mapper [35].

### 2.2. Bioinformatics analysis of thermophilic putative PhaC sequences

A script was written in Python to analyze PhaC sequences for the presence of a lipase(-like) box (https://github.com/MICR-VUB/PhaC-Discovery-Pipeline.git). After screening, protein sequences were categorized as i) harboring a lipase-like box (*lipase-like box*), ii) harboring a lipase box (*lipase box*) or iii) lacking a lipase box (*no lipase box*). This filtering was done based on the presence or absence of a *lipase-like box* ([GS]-X-C-X-[GA]-G) in the protein sequence or a *true lipase box* (G-X-S-X-GG) To this end, a custom pattern recognition script was created in Python. Multiple sequence alignments of *lipase-like box* sequences were visualized using the pyMSAviz package in Python (https://pypi.org/project/pymsaviz/). Genomic environments and genetic synteny between genomes were screened using the Prokaryotic Synteny & Taxonomy Explorer (SyntTax) tool (https://archaea.i2bc.paris-saclay.fr/synttax/) [36]. Finally, phylogenetic trees were created with the Biopython package in Python (https://biopython.org/wiki/Phylo).

### 2.3. Cultivation of bacterial strains

Bacterial strains *T. thermophilus* HB8 DSM 579 and *Paracoccus denitrificans* DSM 413 were obtained from the German Collection of Microorganisms and Cell Cultures (DSMZ). *T. thermophilus* HB27 DSM 7039 was a kind gift from Martine Roovers (research institute LABIRIS).

All bacterial strains were initially incubated overnight in liquid Lysogeny Broth (LB) medium [37,38] while shaking at 75°C, 65°C and 30°C for *T. thermophilus* HB8, *T. thermophilus* HB27 and *P. denitrificans*, respectively. Subsequently, 3% of these precultures were used to inoculate cultures in a mineral salt medium (MSM) to promote PHA production. MSM was prepared according to Pantazaki et al. [18] with some alternations. The medium was obtained by dissolving 15 g carbon source, either sodium gluconate or sucrose, or by adding 15 mL of 10 mM butyric acid or 15 mM octanoic acid as a carbon source, as indicated, with 10 g NaCl, 1.2 g KH_2_PO_4_, 0.26 g K_2_HPO_4_.3H_2_0, 19.9 mg CaCl_2_.2H_2_O, 123 mg MgSO_4_.7H_2_O, 1 g NH_4_Cl and 6 mL of a mineral solution containing (1.96 g H_3_PO_4_, 56 mg FeSO_4_.7H_2_O, 29 mg ZnSO_4_.7H_2_O, 16.7 mg MnSO_4_.H_2_O, 2.5 mg CuSO_4_.5H_2_O, 3 mg Co(NO_3_)_2_.6H_2_O and 6 mg H_3_BO_3_ per liter) in 1 L distilled water. The pH of the medium was adjusted to 7 with NaOH. All cultures were cultivated for 72 hours, upon which cells were harvested by centrifugation at 6000 x *g* for 10 minutes.

### 2.4. Extraction of lipophilic compounds

Two different extraction methods, typically employed for PHA extraction, were conducted to assess the ability of *T. thermophilus* for PHA accumulation in comparison to the prototypical PHA producer *P. denitrificans*.

The first method was performed according to Giedraitytė and Kalėdienė [39] as follows: bacterial strains were cultivated in 30 mL MSM medium, after which cells were collected by centrifugation at 6000 x *g* for 10 minutes. Next, pellets were washed twice with distilled water and resuspended in 10 mL 50 mM sodium phosphate buffer (pH 7). Subsequently, cells were disrupted by an ultrasonic treatment with a Vibra Cell Bioblock Scientific 75043 sonicator during 6 minutes at an amplitude of 20%. Supernatants were removed by centrifugation at 5000 x *g* for 15 minutes and pellets were washed with a 1:1 ethanol:acetone mixture during 1 hour. Hereafter, mixtures containing the washed pellets were each dropped in boiling chloroform. Upon evaporating the solvent, and when the mixture was sufficiently concentrated, the solution was dropped into 10 volumes of 95% ethanol, thereby enabling precipitation. Finally, the obtained lipophilic compounds were separated by centrifugation at 8000 x *g* for 20 minutes and left to dry at room temperature.

The second extraction method was performed as described by Mozejko-Ciesielska et al. [40]. Cell pellets were collected through centrifugation (6000 x *g* for 10 minutes) and snap-frozen in liquid nitrogen. When frozen, the cells were lyophilized for 16 hours in a benchtop lyophilizer (Labconco). This extraction was performed by shaking lyophilized cells, obtained from 100 mL of culture, in 10 mL chloroform at 50°C for 3 hours. This mixture was then filtered through No. 1 Whatman filter paper, followed by precipitation of lipophilic compounds by adding 4 volumes of a chilled 70% methanol solution to the filtrate and leaving it to dry at room temperature.

### 2.5. Gas chromatography

In order to prepare samples for gas chromatography coupled to electron impact mass spectrometry (GC-MS), 2 mg of extracted lipophilic compounds were resuspended in 600 μL of 10% sulfuric acid in methanol, followed by an incubation at 100°C for 4 hours, enabling conversion into β-hydroxycarboxylic acid methyl esters. After allowing the methanol solution to cool to room temperature, 1 mL distilled water and 2 times 0.5 mL dichloromethane were added to the sample, which was shaken vigorously during 1 minute. After phase separation, the organic layers were collected and the solvent was evaporated. Next, *N,O*-Bis(trimethylsilyl)acetamide was added and the mixture was heated to 60°C for 1 hour. This step was performed to obtain the silylated equivalents. Finally, 10 μL of the mixture was diluted with 1 mL ethyl acetate, of which 1 μL was subjected to GC-MS analysis [41]. Resulting methyl esters were analyzed on a Shimadzu GC-MS QP5050A system running with helium as carrier gas and equipped with a split/splitless injector and an Agilent HP-5MS column (column length: 30 m, column diameter: 0.25 mm, film thickness: 0.25 μm). The program used was 2 minutes hold time at 50°C, followed by 15°C/minute to 300°C.

Gas chromatography in combination with a flame ionization detector (GC-FID) was performed directly on biomass. Eight to 11 mg dry biomass was weighed into a crimp vial, followed by the addition of 1 mL of chloroform and 0.8 mL of an esterification mixture with an internal standard (15 % sulphuric acid in methanol with 5 mg/ml benzoic acid as internal standard). This vial was then hermetically closed and incubated at 94°C during 3 hours for esterification. After esterification, the sample was neutralized by transferring the contents of the vial into 0.5 mL of 50 mM NaOH. After shaking and stabilization of the phases, 50 µL was removed from the lower chloroform phase and added to 900 µL of isopropanol in a clean screw-cap vial. This sample was then subjected to GC-FID with a Thermo Scientific Trace 1300 and a Stabilwax, Restek column (30 m; 0.32 mm ID; 0.5 µm DF). Nitrogen was used as carrier gas at a flow rate of 2 mL/min.

### 2.6. Nuclear magnetic resonance spectroscopy

As a means to analyze the extracted compounds with ^1^H nuclear magnetic resonance (NMR) spectroscopy, about 3 mg was solubilized in 0.7 mL deuterated chloroform (CDCl_3_). ^1^H NMR spectra were recorded on a Bruker Avance 250 spectrometer at 250 MHz using a standard pulse sequence. ^1^H NMR chemical shifts (δ) are reported in ppm relative to tetramethylsilane (TMS) and referenced to the residual solvent signal (CDCl_3_: 7.26 ppm).

### 2.7. Fourier Transform Infrared spectroscopy

Fourier Transform Infrared (FTIR) spectroscopy was performed by recording Infrared (IR) spectra using a Nicolet 6700 FTIR spectrophotometer (Thermo Fisher Scientific), operated in a single bounce Attenuated Total Refractance (ATR) mode using the Smart iTR accessory. A diamond plate with a 42° angle of incidence was used. Using this system, 32 scans (resolution 4 cm^-1^) were taken between 600 to 4000 cm^-1^ to form the final IR spectra. FTIR analysis was also applied to poly(3-hydroxybutyrate-*co*-3-hydroxyvalerate) (PHBV, 2% 3-HV) as a reference (Sigma-Aldrich). Spectra were selected between 1900 and 700 cm^-1^ and scaled to a maximum absorbance of 1 within this region.

### 2.8. Differential scanning calorimetry

The thermal properties of the extracted compounds, as well as of PHBV (2% 3-HV) (Goodfellow) as a standard, were analyzed by differential scanning calorimetry (DSC), which was performed using a Discovery DSC (TA Instruments) equipped with a refrigerated cooling system (RCS) under a nitrogen flow (50 mL/min). For each DSC measurement, approximately 3 mg of sample was placed in Tzero pans with Tzero hermetic lids (TA Instruments). Each sample was first heated (20 K/min) to 200°C and then maintained for 1 minute, cooled (20 K/min) to -50°C and maintained for 1 minute and finally increased again (20 K/min) to 200°C and maintained for 1 minute. The melting temperature (*T*_m_) was determined during the second heating cycle, at the maximum of the endothermic peak in het heat flow curve. The glass transition temperature (*T*_g_) was determined as the midpoint of the characteristic change in the baseline of the heat flow curve during the second heating step.

## 3. Results and discussion

### 3.1. Construction of a bioinformatics pipeline to identify PHA producers among thermophiles

To address the gap of a scarcity of examples of extreme thermophilic and hyperthermophilic microbes that produce PHAs, we performed a dedicated screening effort for presence of PHA synthesis-relevant genes in a wide range of thermophilic genomes using a custom-made bioinformatics pipeline (**Figure 1**). To this end, the key enzyme PhaC was selected as an indicative marker for PHA synthesis: in contrast to other PHA synthesis enzymes, PhaC does not harbor homology to enzymes involved in other carbon metabolism pathways and in contrast to (*R*)-3-hydroxyacyl-ACP:CoA transacylase (PhaG) and (*R*)-specific enoyl-CoA hydratase (PhaJ), PhaC is part of all PHA synthesis pathways, not only *mcl* PHA pathways. Phasins (*e.g.* PhaP) and regulators (*e.g.* PhaR) were not considered because of their heterogenous nature [42].

**Figure 1.**
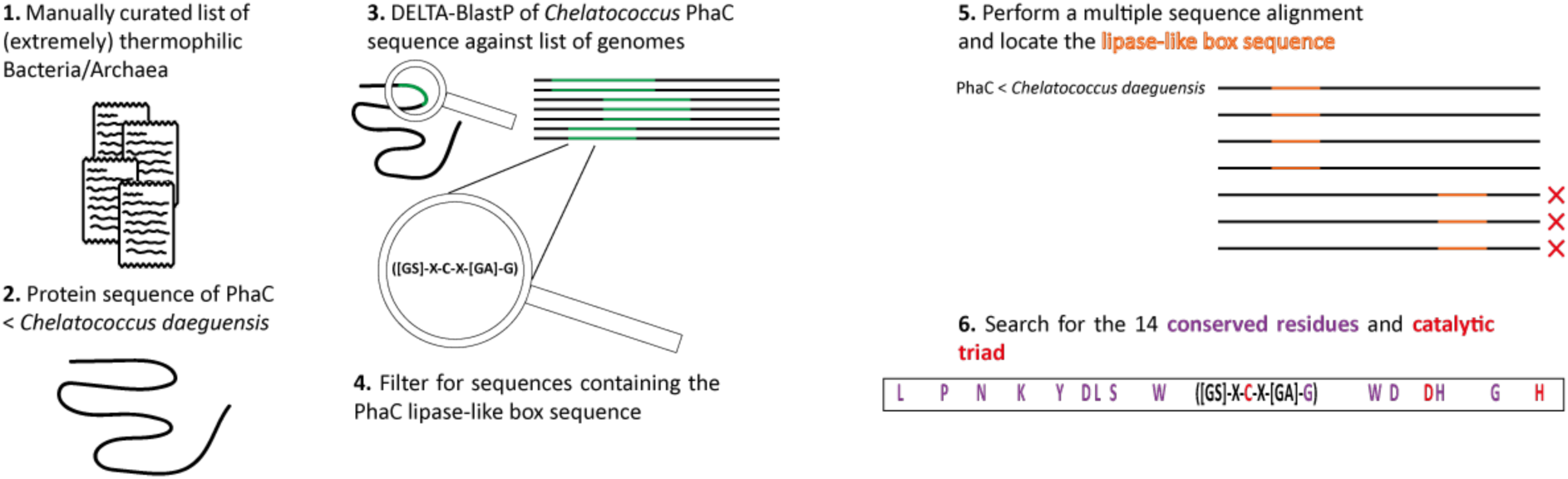
Workflow of the bioinformatics pipeline to find *bona fide* PhaC-encoding genes in the genomes of thermophilic microbial genera.

From a dataset of PhaC sequences covering all enzyme classes (**Supplementary Table S1**), PhaC of *C. daeguensis* TAD1 was selected as a good representative for all four PhaC classes. In general, the core sequence and conserved residues are well conserved over the different classes (**Supplementary Figure S1**). However, we decided to go ahead with a PhaC class I sequence as it is the most general sequence and does not contain an extended C- or N-terminus, thereby it possesses only the core features of a PhaC enzyme. The choice for *C. daeguensis* TAD1 is supported by its high sequence identity to other PhaC sequences (**Supplementary Figure S2**) and it was therefore used as a query in subsequent genome mining. All four categories (moderate thermophilic archaea, bacteria and extreme thermophilic archaea/bacteria) were subjected to a DELTA-BLAST analysis separately (**Figure 1**, **steps 1-3**). DELTA-BLAST was chosen because the structure and domain architecture of enzymes is typically conserved over their amino acid sequence, especially when comparing mesophilic with thermophilic homologs [43]. Two result types emerged: the first type consisted of automatically or manually annotated PhaC subunits. A second type of hits included protein sequences annotated as α/β fold hydrolases, which is the superfamily to which PhaC enzymes belong. The α/β fold hydrolase superfamily is a large family, for example containing all esterases and lipases [44,45].

True PhaC enzymes can be distinguished from other α/β fold hydrolase superfamily members because of the presence of a highly conserved lipase-like box sequence motif ([GS]-X-C-X-[GA]-G). Other members contain a true lipase box sequence motif (G-X-S-X-GG) [46]. Therefore, in a next phase we screened all sequences for the presence of a lipase-like box sequence motif (**Figure 1**, **step 4**). A multiple sequence alignment was performed (**Figure 1**, **step 5**), which revealed that in cases in which a sequence possesses both a lipase and a lipase-like box, the latter was consistently located in a different position as compared to the reference PhaC sequence. A multiple sequence alignment allowed us to visually filter out such hits.

Finally, the remaining hits that passed through the final filtration step were examined to confirm the presence of the essential catalytic triad and, most of, the 14 hyperconserved residues identified during multiple sequence alignments with known PhaC sequences, along with a verification of the essential catalytic triad (**Figure 1**, **step 6**) (**Supplementary Figure S2**).

### 3.2. PhaC occurrence in moderately thermophilic bacteria

Upon screening 8,707 proteomes of moderately thermophilic bacteria, a total of 4,235 lipase box sequences were identified among the top 5,000 hits generated by DELTA-Blast (**Supplementary File S2**). Within this set, 366 sequences featured a lipase-like box, of which 346 sequences displayed a well-aligned position of the box with that of *C. daeguensis* PhaC for 346 sequences (**Supplementary File S3**). Upon further investigation of the catalytic triad, 14 sequences were excluded as they did not contain this essential feature. Finally, screening for the presence of the 14 conserved residues was applied to the remaining sequences. The appearance of the conserved residues ranged between 80 and 97% depending on the residue. This yielded a final list of 332 sequences identified in 142 different species, found on 8,707 proteomes which represents a PhaC retrieval rate of 3.81% that harbor all or most of the conserved residues, including a lipase-like box and the catalytic triad, whilst coming from moderate thermophilic microorganisms (**Supplementary Table S2**). Duplications of PhaCs in some of the detected species might be caused, besides real copies of PhaC, also by sequencing errors and genome misassemblies, especially in draft genomes. Nevertheless, we still observed a well-distributed presence of potential PhaC sequences and, consequently, PHA producers across various bacterial classes. This distribution highlights the widespread nature of this trait among moderately thermophilic bacteria.

**Table 1.**
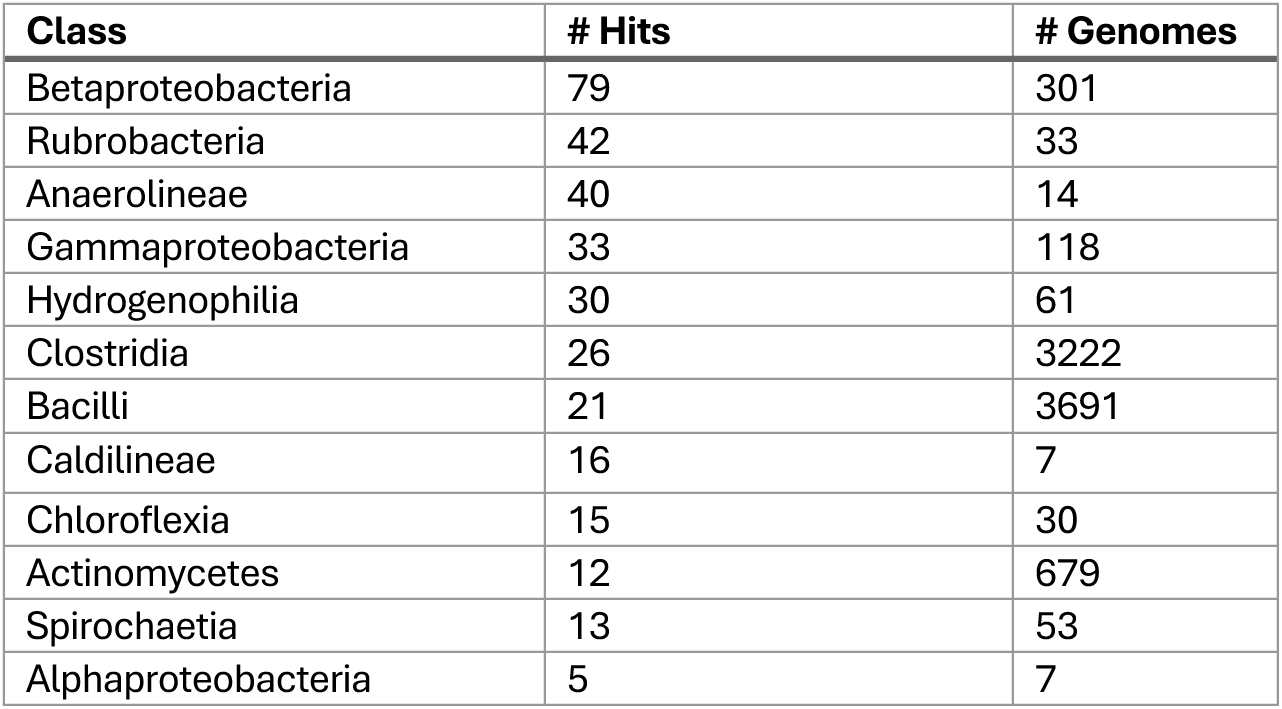
List of the classes of moderately thermophilic microorganisms in which putative PhaC hits were found with the number of screened genomes and hits indicated. A full list of PHA-producing moderate thermophilic bacterial species can be found in **Supplementary Table S2**.

Most of the observed hits originated from species belonging to the class of Actinobacteria and to the genus *Rubrobacter*. Indeed, several *Rubrobacter sp.* have been phenotypically and genotypically identified as PHA producers, for example *Rubrobacter xylanophilus* (T_opt_ = 60°C) [47], which was also detected as a hit in our analysis. Other hits were found in the class of Betaproteobacteria with hits such as *Caldimonas thermodepolymerans* and *Tepidimonas taiwanensis*, which have a T_opt_ between 55°C and 60°C and were also previously identified as PHA producers [48,49]. Finally, we also retrieved sequences that were annotated as α/β fold hydrolases, but seemed to represent novel PhaC sequences in microorganisms which were not described as PHA producers before. An example of this is α/β fold hydrolase WP_033289844.1 of *Amycolatopsis jejuensis*.

### 3.3. PhaC occurrence in moderately thermophilic archaea

Upon investigating proteomes of moderately thermophilic archaea, we obtained a list of 256 sequences harboring a lipase-like box indicative of PhaC. Upon aligning these sequences with known PhaC sequences (**Supplementary File S4**), for 74 of these, the predicted lipase-like boxes aligned with those of the known PhaC sequences. Of these, 9 sequences lack the catalytic triad (**Supplementary File S5**). The remaining 65 sequences, out of a total of 8707 screened genomes (0.76%), can be considered to represent true PhaC sequences, as they harbor a lipase-like box, a complete catalytic triad and the majority of the other conserved residues (ranging between 85-100% depending on the residue) (**Supplementary Table S3**). Again, we have to take into account that some putative PhaC sequences could be duplications in the database, but these seem to be very limited here.

Interestingly, all 65 hits originated from the class Halobacteria spread over 42 species. Several members of Halobacteria were already known for their PHA-producing capabilities, such as the model species *Haloferax mediterranei* [50,51]. Halobacteria typically have an optimal growth temperature between 42°C and 45°C and are thus situated at the lower border of the temperature scale for moderate thermophiles. In contrast, for 743 genomes from 4 other archaeal phylogenetic classes, including Methanomicrobia, Methanobacteria, Thermoplasmata and Methanococci, no PhaC-encoding genes were retrieved. The discovery that all hits were retrieved in a single phylogenetic class contrasts the result for bacterial moderate thermophiles, where hits were spread over 12 different classes.

### 3.4. PhaC occurrence in extreme thermophilic bacteria

Among extreme thermophilic bacterial genera, 164 sequences were identified containing a lipase-like box, within 96 genomes of 834 genomes screened in total (**Supplementary File S6**). None of these hits were annotated as PHA synthases but instead, the majority was annotated as either a homoserine O-acetyltransferase or a α/β hydrolases, which possess a catalytically active serine instead of a cysteine, thus forming a true lipase box [52]. Upon thorough examination of these hits using a multiple sequence alignment (**Supplementary File S7**), we discovered that most hits were predicted to have two sequence motifs: a true lipase box on a position aligned with the lipase-like boxes of confirmed PhaC sequences and a lipase-like box situated in an entirely different location (**Figure 2**).

**Figure 2.**
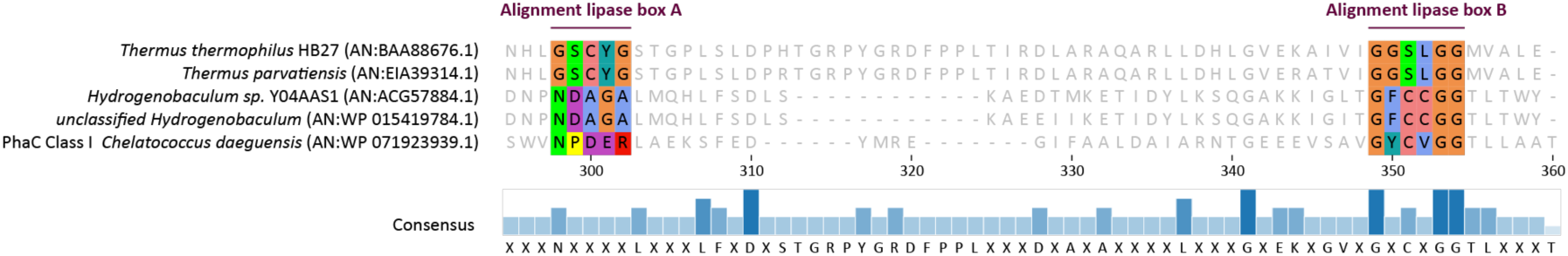
Part of a multiple sequence alignment where the two locations of the lipase boxes are visible. Alignment lipase box A shows where the ‘false’ lipase-like boxes of the *Thermus* sequences align while alignment lipase box B shows where the *Hydrogenobaculum* sequences align with the true lipase-like box as found in *Chelatococcus daeguensis*.

31 sequences were identified that harbor a lipase-like box aligning perfectly with that of known PhaC sequences. Subsequently, these sequences were used to construct a new multiple sequence alignment to ensure the presence of the 12 highly conserved amino acid residues. Similar to moderate thermophiles, we applied additional filtering based on the presence of the other two catalytic residues of the catalytic triad. Among the hits from extreme bacterial thermophiles, only one sequence harbours the complete catalytic triad essential for PHA synthase. This sequence, with accession number WP_0154186861.1, is a multispecies entry which comprises two proteins annotated as a carboxylesterase (ACG56724.1 and PMP61664.1 from *Hydrogenobaculum sp.* Y04AAS1 and an unclassified *Hydrogenobaculum sp.* respectively). Although they are interesting as putative extremely thermophilic PHA synthases, these sequences appeared only distantly related to true PhaC sequences as none of the conserved residues were present. Moreover, these putative hits were located in the genome of *Hydrogenobaculum* sp. Y04AAS1 surrounded by non-PHA-related genes (**Figure 3**), including genes encoding enzymes associated with amino acid metabolism. This suggests an alternative function for these hits, distinct from PHA biosynthesis. Thus, no hits were found for bacterial extreme thermophiles with the present methods.

**Figure 3.**
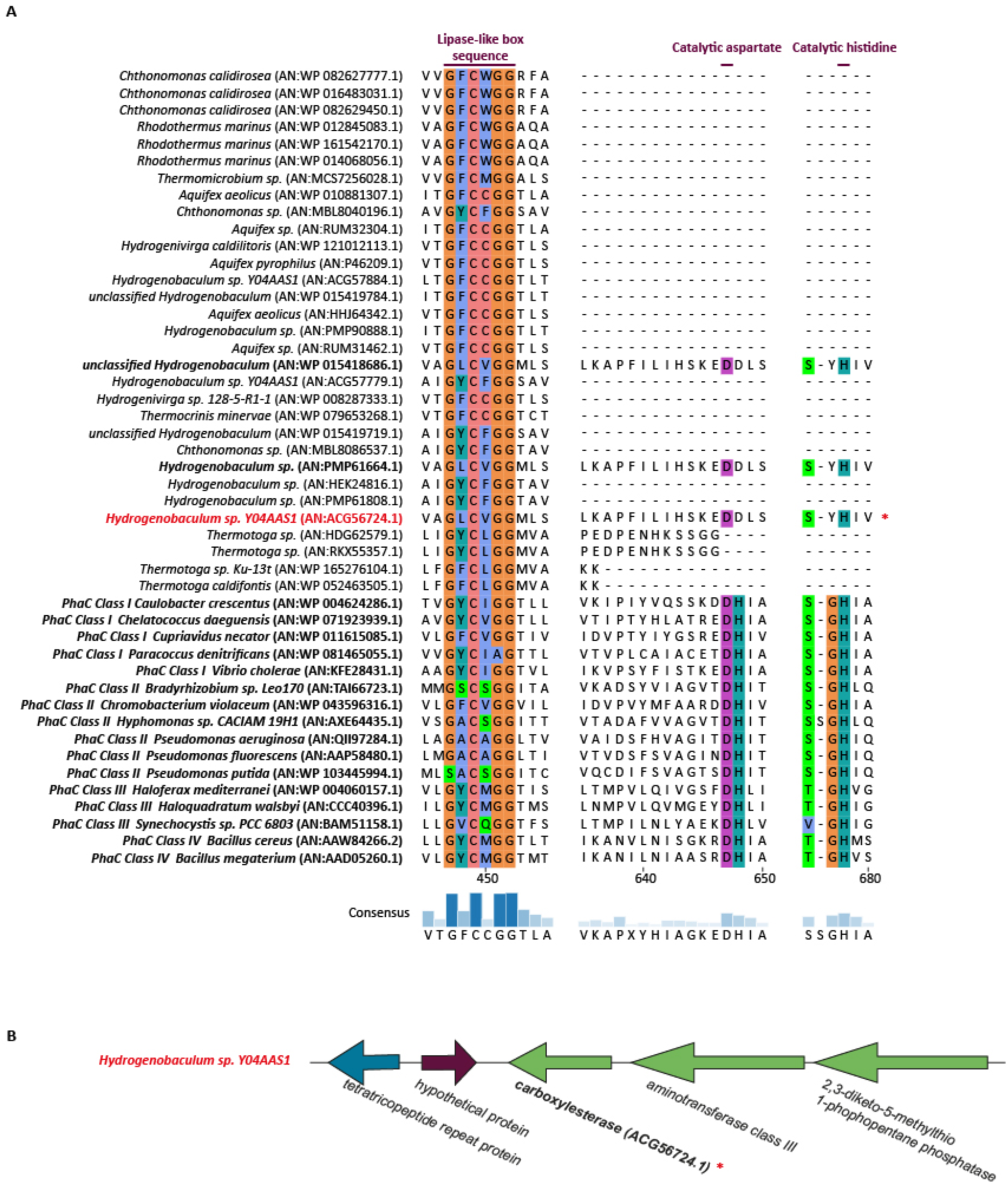
**(A)** Part of a multiple sequence alignment between 16 known and validated PhaC sequences and 31 hits in extreme thermophiles which possess a lipase-like box that aligns perfectly with those of the known PhaC sequences. Sequences marked with a red asterisk are the only sequences which harbour the full catalytic triad. A full multiple sequence alignment can be found in **Supplementary File S7**. **(B)** Genomic organisation of one of the hits, marked in bold, found in *Hydrogenobaculum* sp.

It was surprising that no hits within the *Geobacillus* genera were found, as they were previously proposed as candidates for extremely thermophilic PHA production processes [53]. Similarly, we did not detect the PHA synthase among the members of the genus *Thermus*, despite prior reports suggesting that *Thermus thermophilus* is an extreme thermophilic PHA producer [18,19,20].

### 3.5. Prediction of PhaC occurrence in extreme thermophilic archaea

The difference between hits in moderate thermophiles and extreme thermophiles was even more pronounced within archaeal genera, in which only 26 hits were found (**Supplementary File S8**). Similar to the findings in bacterial genera, no sequences were annotated as PHA synthases, but only as α/β fold hydrolase proteins. Among these proteins, 19 harbor a lipase-like box that aligns perfectly with that of known PhaC sequences. Interestingly, 7 of these hits, out of a total of 860 genomes screened (0.8%), possess the complete catalytic triad and all 12 conserved residues (**Figure 4**) (**Supplementary File S9**). Furthermore, the construction of a phylogenetic tree using the neighbor-joining algorithm clearly showed that these seven sequences formed a distinct cluster with class III and class IV PhaC sequences, while the other 19 hits lacking the twelve conserved residues formed a separate outgroup (**Figure 5**).

**Figure 4.**
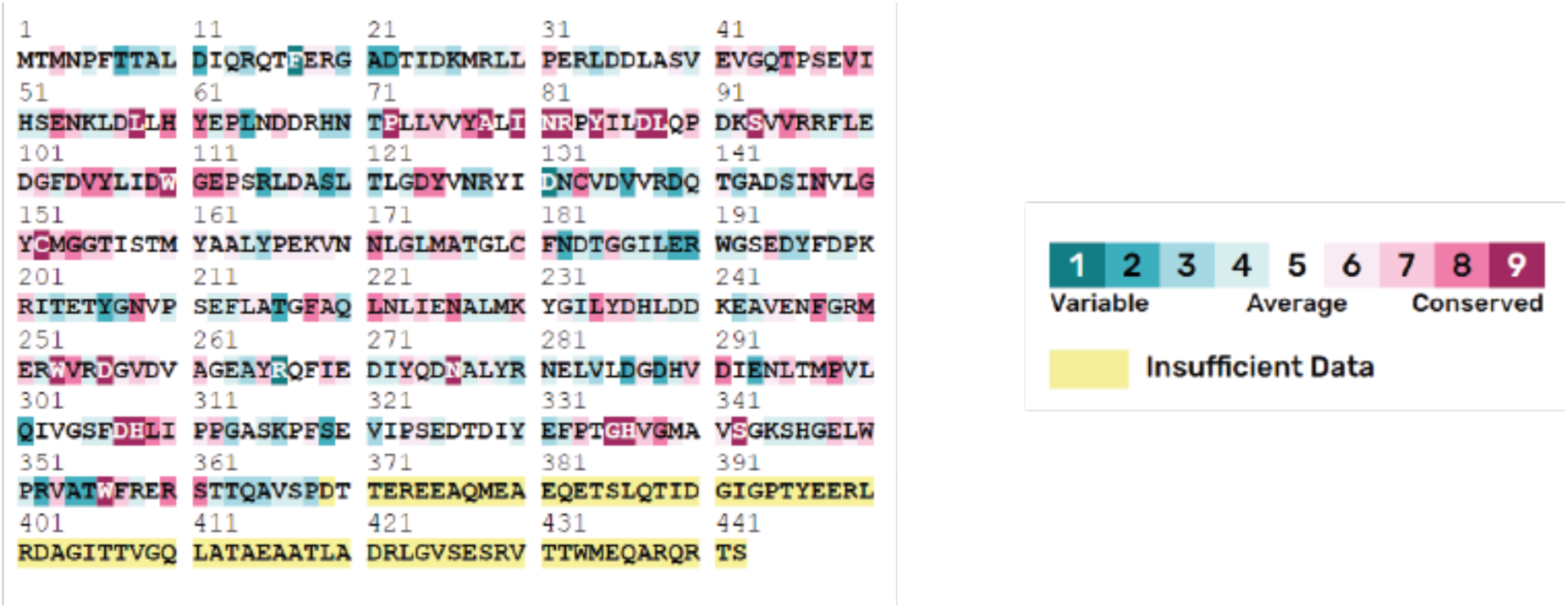
Consensus sequence of the positive putative PhaC hits found within archaeal extreme thermophiles and the known PhaC sequences. Residues that were present in all queries of the multiple sequence alignments are found boxed in purple with a white bold indication of the one-letter amino acid abbreviation of the residue. Here, all 12 conserved residues are indeed present in all sequences. This scheme was created with the ConSurf web server.

**Figure 5.**
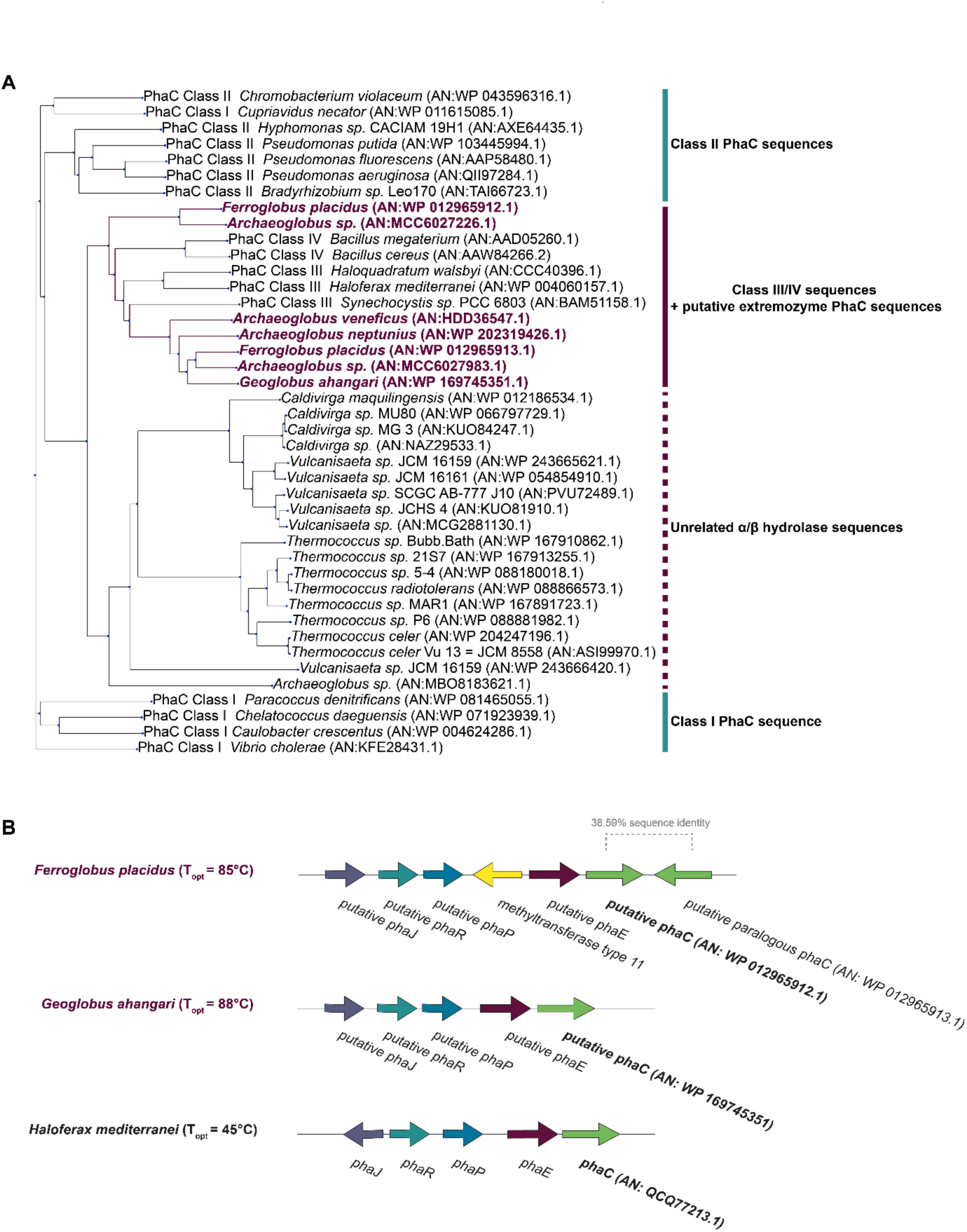
**(A)** Phylogenetic tree of all hits that were obtained upon screening for the presence of PhaC sequences in the genomes of extremely thermophilic archaea. Microorganisms and accession codes in purple represent the seven hits which were found to be true PhaC sequences and cluster together with other class III and class IV PhaC sequences. All the other hits cluster together in a group that we indicated with alpha/beta hydrolase sequences. (**B)** Genomic organization of two of the PhaC hits found in *F. placidus* and *G. ahangari*. Both show a high degree of similarity to the genomic organization of the PHA genes in *H. mediterranei*, which is an archaeal model organism for archaeal PHA biosynthesis.

All seven sequences belonged to the *Ferroglobus*, *Archaeoglobus* or *Geoglobus* genera, and all of them belong to the family Archaeoglobaceae **(Supplementary Table S4).** These genera are hyperthermophilic, strictly anaerobic species originally isolated from marine hydrothermal systems [54]. Examining the genomic neighborhood of the hits in *Ferroglobus placidus* (T_opt_ = 85°C) and *Archaeoglobus ahangari* (T_opt_ = 88°C), which have fully sequenced genomes, showed a gene organization similar to that of the *pha* genes in *Haloferax mediterranei* (T_opt_ = 45°C), where a smaller gene upstream of the hits could potentially be the *phaE* subunit gene. Moreover, homologs of the PHA-regulator *phaR* and phasin gene *phaP*, were found near the hit. These could be arranged in the typical *phaRP* operon and have a 96% chance of being an operon according to operon_mapper. Although the presence of a *phaR*-like regulator in *Ferroglobus* sp. was observed previously, it was unknown that it clustered within a genomic region bearing several PHA-related genes [55].

The genomic organization of the PHA gene cluster in these archaea was observed to resemble the one from *Haloferax mediterranei*, which is not unexpected as the genera *Haloferax*, *Geoglobus* and *Ferroglobus* all belong to the phylum Euryarchaeota. However, it is surprising that the hits in extreme thermophiles were so scarce and clustered in only a few genera, once again as observed with moderate thermophilic archaea. Furthermore, these hits demonstrate the effectiveness of our bioinformatics pipeline in picking up sequences within extreme thermophiles and that, if classical PhaC enzymes were present in the genomes of bacterial extreme thermophiles, our pipeline would likely have detected them. This study marks the first genotypic characterization of an archaeal extreme thermophile in which PHA biosynthesis genes are identified.

3.6. Experimental investigation of the ability of *Thermus thermophilus* to accumulate polyhydroxyalkanoates

The absence of *phaC* genes in extreme bacterial thermophiles such as *Thermus* contradicts earlier findings suggesting that *Thermus thermophilus* HB8 is a PHA producer [18]. To address this contradiction, two strains of *T. thermophilus*, namely HB8 and HB27, were selected for an in-depth experimental exploration of their PHA production capabilities. Different carbon sources were used in combination with MSM that was reported in previous literature to enhance PHA accumulation [18]. The use of sodium octanoate as a sole carbon source did not result in any growth of both strains, while sodium gluconate supported the growth by only a low growth rate. These results are contradictory to what was previously mentioned [20]. For the *T. thermophilus* cultures cultivated on sodium gluconate, as well as a *Paracoccus denitrificans* culture as a positive control, two different PHA extraction protocols were performed [39,40]. In both cases, the obtained powder from *T. thermophilus* cells displayed a brownish appearance, which is distinct from the white color typically observed for extracted PHA, as was also the case for the *P. denitrificans* extract (**Figure 6**).

**Figure 6.**
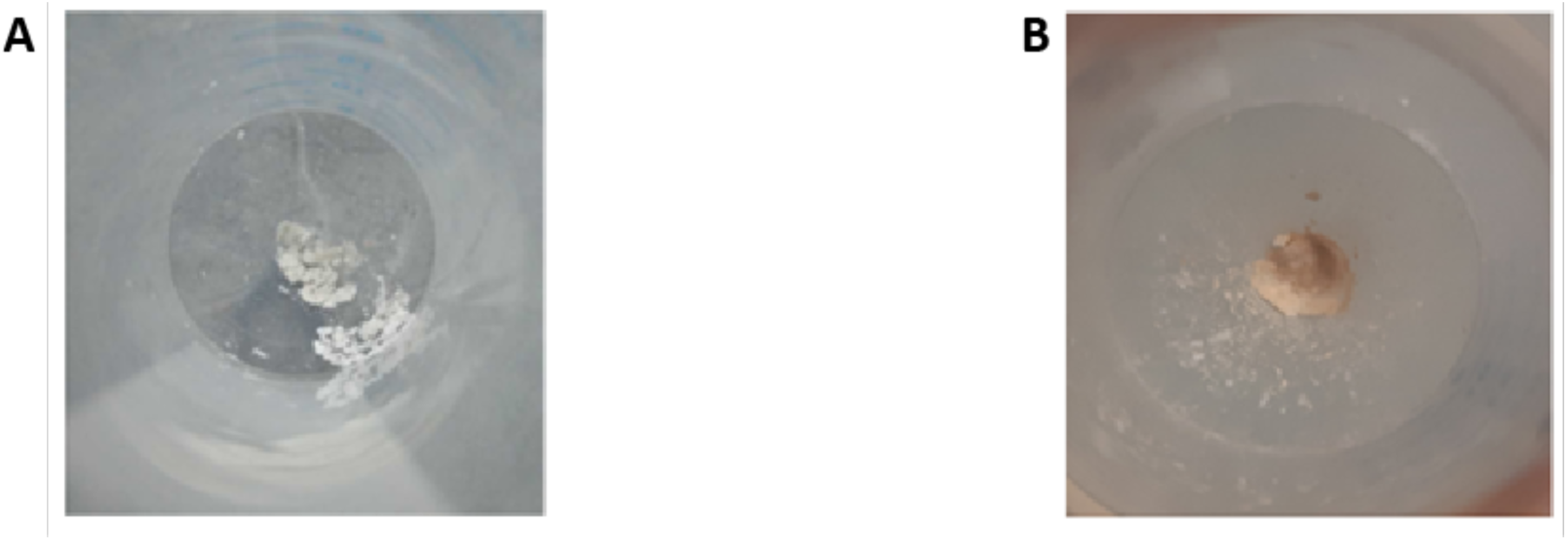
Powder extracts obtained after applying a PHA extraction protocol (sonication-chloroform method) on *P. denitrificans* **(A)** and *T. thermophilus* **(B)**.

The samples were subsequently chemically analyzed using different techniques, employing a reference commercial standard of polypoly(3-hydroxybutyrate-co-3-hydroxyvalerate). *T. thermophilus* HB8 and HB27 extracts were both examined in the chemical analysis and resulted in similar results. FTIR confirmed the complete absence of PHA polymers *T. thermophilus* and its presence in the *P. denitrificans* sample (**Figure 7**). Indeed, the spectrum corresponding to the *P. denitrificans* displays characteristic peaks for PHA polymers. More specifically, the peaks located at 1719 cm^-1^ (C=O stretch), 1453 cm^-1^ (CH_2_ aliphatic stretching), 1379 cm^-1^ (CH_3_ vibration), 1277 and 1228 cm^-1^ (C-O stretching), are indicative of PHAs.

**Figure 7.**
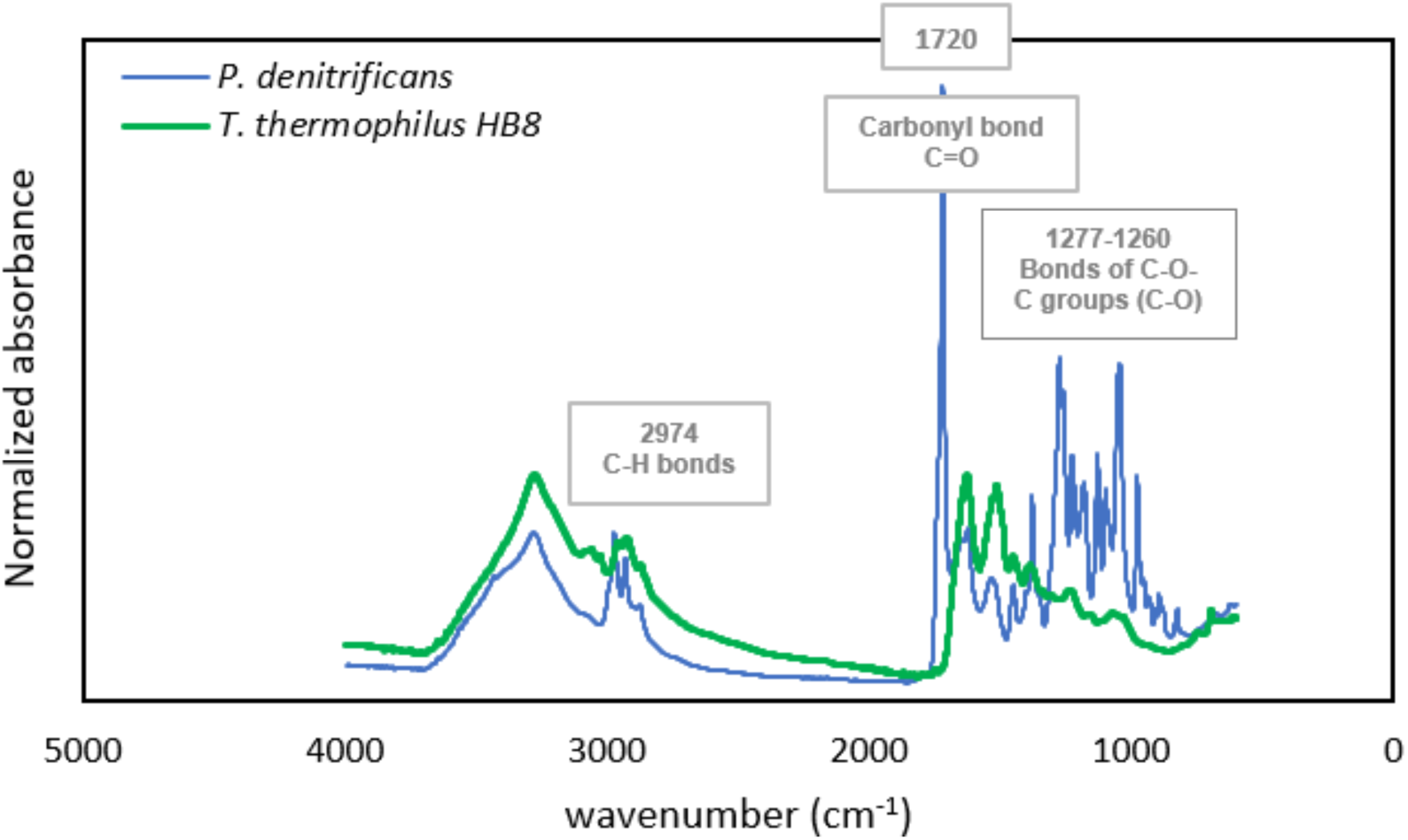
FTIR spectrum of extracts from *P. denitrificans* and *T. thermophilus* HB8 cells grown on MSM medium. Characteristic PHA peaks in the *P. denitrificans* spectrum are absent in the spectrum from the *T. thermophilus* HB8 extract.

The absence of PHA in the *T. thermophilus* samples was also confirmed by GC-MS (**Supplementary Figure S3A** and **C**), in which a main single characteristic peak with a retention time of 6.2 minutes was only revealed in the *P. denitrificans* sample, but absent in the *T. thermophilus* sample. Upon analysis of this characteristic peak by mass spectrometry (**Supplementary Figure S3B**), it was shown to be the typical fingerprint of the 3-(trimethylsilyl)-methyl ester of 3-hydroxybutanoate, the silylated derivative of the monomer of poly(3-hydroxybuyrate). In contrast, small peaks were noticed in the chromatogram of the *T. thermophilus* sample with retention time starting from 7.39 minutes. Upon analysis of these peaks by mass spectrometry, the resulting fingerprints revealed that these represent products of side reactions of the silylating agent used in sample preparation. A further characterization of the extracted material was done using differential scanning calorimetry which also showed a difference between the *P. denitrificans* sample and the *T. thermophilus* sample as there was no sign of any thermal transition in the latter (**Supplementary File S4**).

^1^H NMR was used to chemically analyze the extracted samples from both strains. Also in this case, characteristic peaks associated with PHB were distinguished in the *P. denitrificans* sample while being absent in the extracted sample from *T. thermophilus* (**Supplementary Figure S5**). The largest peak around 1.25 ppm was attributed to protons of the methyl group which is the side chain of poly(3-hydroxybutyrate). The two peaks found at 2.5 ppm formed a double doublet and were due to the protons of CH_2_ in the backbone of the polymer. The peak associated with the PHB structure, and the highest chemical shift (5.2 ppm) was attributed to the proton of the carbon that was bound to the carboxyl group.

Finally, we also attempted to detect PHA in cells of the strains belonging to the genus *Thermus* using a routine technique for PHA identification and quantification—direct GC-FID analysis of the dried biomass. However, we did not observe any traces of PHA polymers in any of the samples analyzed (data not shown). It should be noted that at least one other research group has also been unable to identify PHA synthesis capabilities in *Thermus thermophilus* at either the genomic or phenotypic level [56].

### 3.7. Hypothetical mechanisms and reasons for the scarcity of PHA synthesis in extreme thermophiles

We would like to propose several hypothetical explanations for the phenomenon of PHA synthesis being so rare among extreme thermophiles. The first hypothesis is related to the energy metabolism of extreme thermophiles and hyperthermophiles. Given the well-known fact that oxygen solubility decreases with increasing temperature, these microorganisms are likely to prefer anaerobic metabolism, which generally yields less energy than aerobic metabolism. As a result, they may not need to store extensive amounts of energy and carbon resources in the form of PHA granules. While there are known PHA producers among mesophilic anaerobes [57,58], it is highly likely that microorganisms adapted to high temperatures have additional energy demands associated with survival strategies under specific stress conditions, reducing their need and capacity to invest energy into PHA as an energy storage compound.

The second reason why PHA synthesis is very rare among very hot niches accommodating thermophiles could be that presence of PHA granules results has serious impacts on physicochemical properties of cytoplasm which are incompatible with high-temperature conditions. First, the accumulation of PHA granules significantly increases molecular crowding in the bacterial cytoplasm, leading to notable effects on density of intracellular space. Molecular crowding impacts cellular processes by altering the effective concentration, diffusion, and interactions of macromolecules [59]. For example, crowding limits the speed of translation due to the reduced diffusion of bulky tRNA complexes, ultimately constraining cell growth rates [60]. This crowded environment also affects biophysical processes, such as protein folding, aggregation, and the assembly of macromolecular structures, as explained by the excluded volume effect [61,62]. In *Rhodobacter sphaeroides*, variations in crowding, driven by different numbers of intracytoplasmic membrane vesicles, have been shown to alter protein diffusion rates, suggesting that physiological activities can exhibit distinct kinetics under varying growth conditions [63]. Theoretical models and simulations further highlight that crowding significantly impacts the kinetics and thermodynamics of molecular interactions [64].

Molecular crowding also influences the solubility and diffusion of small molecules like oxygen within the cytoplasm. Specifically, the effective diffusion rate of oxygen is reduced in a crowded environment due to a decrease in available volume fraction and long-range interactions between crowders and diffusers [65]. These adverse effects of overcrowding may become critical at higher temperatures, as temperature alters the physical state and interactions of molecules within the cytoplasm, thereby impacting diffusion rates and biochemical reaction kinetics [62].

Furthermore, we have previously reported that in a mesotrophic bacterium, *C. necator*, the presence of PHA granules affects the activity of intracellular water, making it more susceptible to transmembrane transport [5,66]. It is therefore likely that further enhancement of transmembrane water loss at higher temperatures could have detrimental effects, compromising both the structural integrity and physiological functions of the cells. Water loss from bacterial cells, especially in high-osmolarity environments, results in cytoplasmic dehydration and a reduction in turgor pressure, which in extreme cases can lead to plasmolysis and inhibit bacterial growth [67]. This increased dehydration, combined with the above-described effects of molecular crowding, could elevate the effective concentration of cellular constituents beyond the lethal threshold.

Additionally, it can be speculated that extremely high temperatures may be incompatible with the architecture of native PHA granules. The hydrophobic PHA polymer is separated from the hydrophilic cytoplasm by a layer of PHA granule-associated proteins, which are attached to the surface of the granules and serve as a functional interface [68]. It is well known that native PHA granules are formed by the polymer in a thermodynamically unfavourable amorphous state. This state has crucial biophysical and biological implications. Due to their amorphous nature, the viscoelastic properties of native granules resemble a supercooled liquid rather than solid polymer particles, which is an important factor in making PHA granules more compatible within bacterial cells [66]. Furthermore, intracellular PHA depolymerases, enzymes responsible for metabolizing PHA storage materials when external carbon sources are limited, are specific to the amorphous state of the PHA polymer [69]. Therefore, it is essential for bacterial cells to maintain PHA granules in an amorphous state. In fact, the thermodynamically metastable state of PHA granules is synergistically stabilized by two mechanisms: (i) kinetically, by the low rate of crystallization within the limited volume of “small” PHA granules, and (ii) by the presence of water within the granules, which acts as a plasticizer, protecting the polymer from crystallization [70]. It is highly likely that at higher temperatures, as the mobility of PHA polymer chains substantially increases, despite the presence of PHA-granules-associated-proteins PHA granules tend to aggregate, which could have severe consequences for the intracellular architecture of bacterial cells, negatively affecting cell division and other processes. Moreover, due to the increase in granule volume and the enhanced mobility of PHA chains and also intragranular “plasticizing” water, which could consequently be removed from the granules, the polymer has a higher tendency to crystallize. Therefore, the scarcity of PHA granules in extreme thermophiles and hyperthermophiles could also be a consequence of the challenges associated with maintaining PHA granules in a separated and amorphous state at elevated temperatures.

## 4. Conclusions

In conclusion, our investigation indicates that the potential for PHA biosynthesis among prokaryotic organisms is primarily restricted to psychrophiles, mesophiles, and moderate thermophiles, with only a limited number of exceptions. Various hypothetical impediments may elucidate the infrequency of PHA producers within extreme thermophiles and hyperthermophiles. One plausible explanation is that at elevated temperatures, microorganisms may prioritize the allocation of their finite energy resources towards fundamental biological functions, rather than towards energy storage mechanisms. Furthermore, the existence of PHA granules may adversely affect the biophysical characteristics of the cytoplasmic environment, and the structural integrity of native PHA granules could present challenges associated with stability and separation at exceedingly high temperatures.

Conversely, our bioinformatics assessment indicates possible exceptions among extremely thermophilic archaea, particularly within the genera *Ferroglobus*, *Archaeoglobus* and *Geoglobus*. This observation suggests that, although uncommon, PHA biosynthesis at extreme thermal conditions is not entirely impossible. As a result, additional research is warranted to ascertain whether these thermophilic prokaryotes possess the capability for PHA synthesis. If validated, the investigation of their PHA metabolic pathways and the functional attributes of pivotal proteins involved in PHA biosynthesis, degradation, and granule stabilization could provide significant insights. Elucidating the mechanisms that facilitate PHA synthesis under extremely high temperatures could enhance our understanding of extremophiles existing at the limits of life and potentially facilitate biotechnological applications of PHA production in extreme conditions through metabolic engineering and synthetic biology.

## CRediT authorship contribution statement

**Brendan Schroyen**: Conceptualization, Software, Formal analysis, Investigation, Data Curation, Writing – Original Draft, Visualization. **Radwa Moanis**: Conceptualization, Investigation, Writing – Original Draft, Visualization. **Hannelore Geeraert**: Methodology, Writing – Review & Editing. **Niko Van den Brande**: Methodology, Resources, Writing – Review & Editing. **Ulrich Hennecke**: Methodology, Resources, Writing – Review & Editing. **Stanislav Obruča**: Investigation, Writing – Original Draft. **Iva Buchtíková**: Investigation, Writing – Review & Editing. **Karel Sedlář**: Investigation, Writing – Original Draft. **Petr Sedláček**: Investigation, Writing – Review & Editing. **Eveline Peeters**: Conceptualization, Resources, Project administration, Writing – Review & Editing, Funding acquisition.

## Acknowledgments

The authors are grateful to Martine Roovers for the kind gift of the *Thermus thermophilus* strain and to Karl Jonckheere for technical assistance.

## Data availability

All data are available in Supplementary Materials and/or in a Zenodo dataset 10.5281/zenodo.15221919.

