## Supplementary Materials for "Tracking polyhydroxyalkanoate biosynthesis in thermophilic microorganisms"

An accompanying dataset has been published on Zenodo with doi 10.5281/zenodo.15221919.

Supplementary File S1 - List of archaeal/bacterial genera/classes and their classification based on optimal growth temperature.

Supplementary Table S1 - List of microorganisms of which the PhaC sequences were used in the bioinformatics pipeline.

Supplementary Figure S1- Multiple sequence alignment of the 16 well-described or automatically annotated PhaC sequences.

Supplementary Figure S2 - Pairwise percent identity matrix of aligned PhaC protein sequences.

Supplementary File S2 - List of putative PhaC sequences with lipase-like box motifs in bacterial moderate thermophiles. Lines in the list highlighted in bold red contained putative PhaC sequences in which the lipase-like box did not align with the lipase-like box of the reference PhaC sequences and are thus not taken further into the analysis.

Supplementary File S3 (Zenodo) – Full multiple sequence alignment of the putative PhaC sequences found in bacterial moderate thermophiles together with 16 reference PhaC sequences (366 putative PhaC sequences + 16 reference sequences).

Supplementary Table S2 - Filtered list of PhaC sequences in bacterial moderate thermophiles.

Supplementary File S4 - List of putative PhaC sequences with lipase-like box motifs in archaeal moderate thermophiles. Lines in the list highlighted in bold red contained putative PhaC sequences in which the lipase-like box did not align with the lipase-like box of the reference PhaC sequences and are thus not taken further into the analysis.

Supplementary File S5 (Zenodo) - Full multiple sequence alignment of the putative PhaC sequences found in archaeal moderate thermophiles together with 16 reference PhaC sequences (256 putative PhaC sequences + 16 reference sequences).

Supplementary Table S3 - Filtered list of PhaC sequences in archaeal moderate thermophiles.

Supplementary File S6 – List of putative PhaC sequences with lipase-like box motifs in bacterial extreme thermophiles. Lines in the list highlighted in bold red contained putative PhaC sequences in which the lipase-like box did not align with the lipase-like box of the reference PhaC sequences and are thus not taken further into the analysis.

Supplementary File S7 (Zenodo) - Full multiple sequence alignment of the putative PhaC sequences found in bacterial extreme thermophiles together with 16 reference PhaC sequences (164 putative PhaC sequences + 16 reference sequences).

Supplementary File S8 - List of putative PhaC sequences with lipase-like box motifs in archaeal extreme thermophiles. Lines in the list highlighted in bold red contained putative PhaC sequences in which the lipase-like box did not align with the lipase-like box of the reference PhaC sequences and are thus not taken further into the analysis.

Supplementary File S9 (Zenodo) - Full multiple sequence alignment of the putative PhaC sequences found in archaeal extreme thermophiles together with 16 reference PhaC sequences (26 putative PhaC sequences + 16 reference sequences).

Supplementary Table S4 - Filtered list of PhaC sequences in archaeal extreme thermophiles.

Supplementary Figure S3 – Results of the GC-MS analysis.

Supplementary Figure S4 – Results of the DSC analysis.

Supplementary Figure S5 – Results of the $^{1}$H-NMR analysis.

**Supplementary Table S1**: List of microorganisms of which the PhaC sequences were used in the bioinformatics pipeline. Well-described microorganisms and their PhaC sequence are accompanied by their descriptive paper. Whenever an automatic annotation of NCBI was used, AA has been written in the list of sources.

| Microorganism | Accesion number | PhaC Class | Source |
|---|---|---|---|
| *Caulobacter crescentus* | WP_004624286.1 | I | Qi *et al.*, 2001 |
| *Chelatococcus daeguensis* | WP_071923939.1 | I | AA |
| *Cupriavidus necator* | WP_011615085.1 | I | Wittenborn *et al.*, 2016 |
| *Paracoccus denitrificans* | WP_081465055.1 | I | AA |
| *Vibrio cholerae* | KFE28431.1 | I | AA |
| *Bradyrhizobium sp.* Leo170 | TAI66723.1 | II | AA |
| *Chromobacterium violaceum* | WP_043596316.1 | II | AA |
| *Hyphomonas sp.* CACIAM 19H1 | AXE64435.1 | II | AA |
| *Pseudomonas aeruginosa* | QII97284.1 | II | Amara *et al.*, 2003 |
| *Pseudomonas fluorescens* | AAP58480.1 | II | AA |
| *Pseudomonas putida* | WP_103445994.1 | II | AA |
| *Haloferax mediterranei* | WP_004060157.1 | III | Lu *et al.*, 2008 |
| *Haloquadratum walsbyi* | CCC40396.1 | III | Han *et al.*, 2010 |
| *Synechocystis sp.* PCC 6803 | BAM51158.1 | III | Numata *et al.*, 2015 |
| *Bacillus cereus* | AAW84266.2 | IV | Kihara *et al.*, 2017 |
| *Bacillus megaterium* | AAD05260.1 | IV | McCool *et al.*, 2001 |

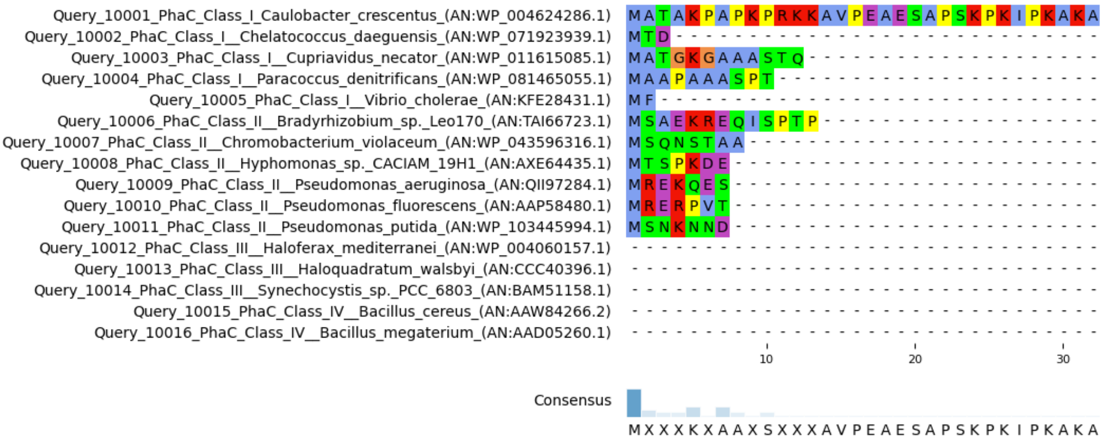

Query_10001_PhaC_Class_I_Caulobacter_crescentus_(AN:WP_004624286.1)
Query_10002_PhaC_Class_I__Chelatococcus_daeguensis_(AN:WP_071923939.1)
Query_10003_PhaC_Class_I__Cupriavidus_necator_(AN:WP_011615085.1)
Query_10004_PhaC_Class_I__Paracoccus_denitrificans_(AN:WP_081465055.1)
Query_10005_PhaC_Class_I__Vibrio_cholerae_(AN:KFE28431.1)
Query_10006_PhaC_Class_II__Bradyrhizobium_sp._Leo170_(AN:TAI66723.1)
Query_10007_PhaC_Class_II__Chromobacterium_violaceum_(AN:WP_043596316.1)
Query_10008_PhaC_Class_II__Hyphomonas_sp._CACIAM_19H1_(AN:AXE64435.1)
Query_10009_PhaC_Class_II__Pseudomonas_aeruginosa_(AN:QII97284.1)
Query_10010_PhaC_Class_II__Pseudomonas_fluorescens_(AN:AAP58480.1)
Query_10011_PhaC_Class_II__Pseudomonas_putida_(AN:WP_103445994.1)
Query_10012_PhaC_Class_III__Haloferax_mediterranei_(AN:WP_004060157.1)
Query_10013_PhaC_Class_III__Haloquadratum_walsbyi_(AN:CCC40396.1)
Query_10014_PhaC_Class_III__Synechocystis_sp._PCC_6803_(AN:BAM51158.1)
Query_10015_PhaC_Class_IV__Bacillus_cereus_(AN:AAW84266.2)
Query_10016_PhaC_Class_IV__Bacillus_megaterium_(AN:AAD05260.1)
10
20
30
Consensus
M X X X K X A A X S X X X A V P E A E S A P S K P K I P K A K A


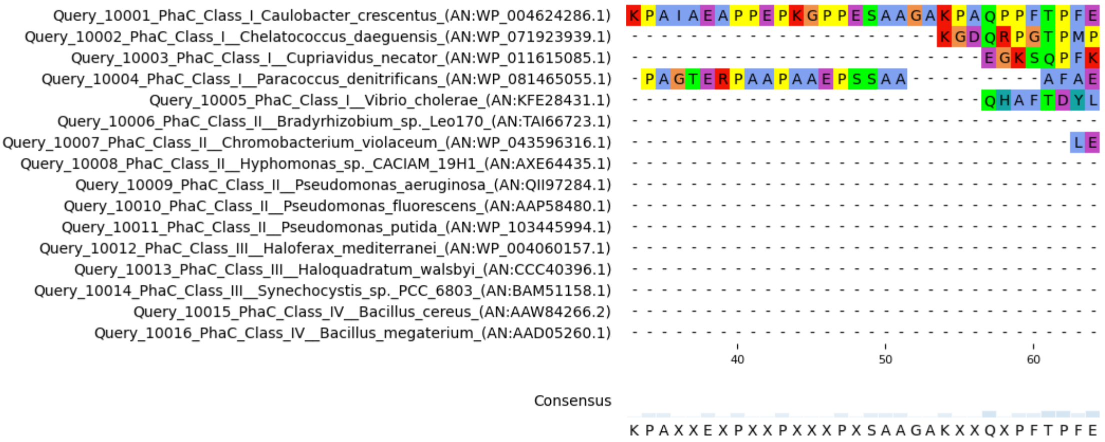

Query_10001_PhaC_Class_I_Caulobacter_crescentus_(AN:WP_004624286.1)
Query_10002_PhaC_Class_I__Chelatococcus_daeguensis_(AN:WP_071923939.1)
Query_10003_PhaC_Class_I__Cupriavidus_necator_(AN:WP_011615085.1)
Query_10004_PhaC_Class_I__Paracoccus_denitrificans_(AN:WP_081465055.1)
Query_10005_PhaC_Class_I__Vibrio_cholerae_(AN:KFE28431.1)
Query_10006_PhaC_Class_II__Bradyrhizobium_sp._Leo170_(AN:TAI66723.1)
Query_10007_PhaC_Class_II__Chromobacterium_violaceum_(AN:WP_043596316.1)
Query_10008_PhaC_Class_II__Hyphomonas_sp._CACIAM_19H1_(AN:AXE64435.1)
Query_10009_PhaC_Class_II__Pseudomonas_aeruginosa_(AN:QII97284.1)
Query_10010_PhaC_Class_II__Pseudomonas_fluorescens_(AN:AAP58480.1)
Query_10011_PhaC_Class_II__Pseudomonas_putida_(AN:WP_103445994.1)
Query_10012_PhaC_Class_III__Haloferax_mediterranei_(AN:WP_004060157.1)
Query_10013_PhaC_Class_III__Haloquadratum_walsbyi_(AN:CCC40396.1)
Query_10014_PhaC_Class_III__Synechocystis_sp._PCC_6803_(AN:BAM51158.1)
Query_10015_PhaC_Class_IV__Bacillus_cereus_(AN:AAW84266.2)
Query_10016_PhaC_Class_IV__Bacillus_megaterium_(AN:AAD05260.1)
40
50
60
Consensus
K P A X X E X P X X P X X X P X S A A G A K X X Q X P F T P F E


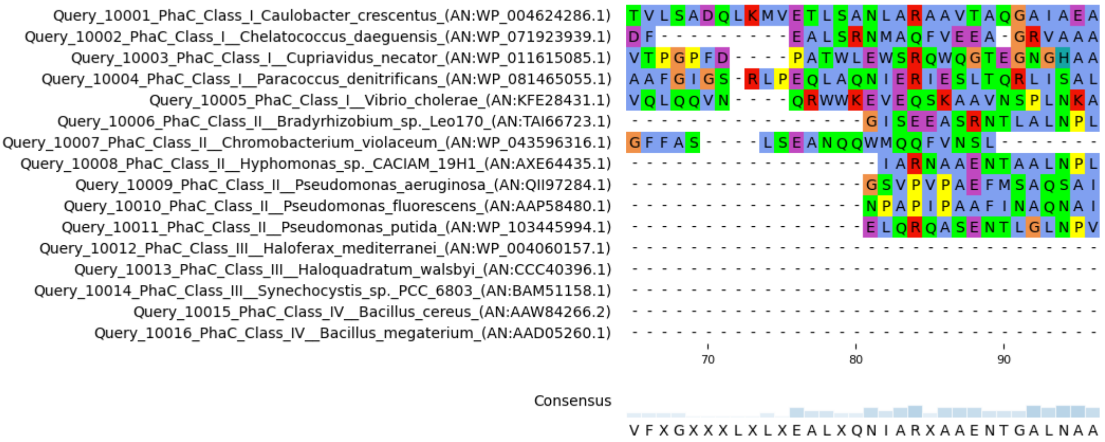

Query_10001_PhaC_Class_I_Caulobacter_crescentus_(AN:WP_004624286.1)
Query_10002_PhaC_Class_I__Chelatococcus_daeguensis_(AN:WP_071923939.1)
Query_10003_PhaC_Class_I__Cupriavidus_necator_(AN:WP_011615085.1)
Query_10004_PhaC_Class_I__Paracoccus_denitrificans_(AN:WP_081465055.1)
Query_10005_PhaC_Class_I__Vibrio_cholerae_(AN:KFE28431.1)
Query_10006_PhaC_Class_II__Bradyrhizobium_sp._Leo170_(AN:TAI66723.1)
Query_10007_PhaC_Class_II__Chromobacterium_violaceum_(AN:WP_043596316.1)
Query_10008_PhaC_Class_II__Hyphomonas_sp._CACIAM_19H1_(AN:AXE64435.1)
Query_10009_PhaC_Class_II__Pseudomonas_aeruginosa_(AN:QII97284.1)
Query_10010_PhaC_Class_II__Pseudomonas_fluorescens_(AN:AAP58480.1)
Query_10011_PhaC_Class_II__Pseudomonas_putida_(AN:WP_103445994.1)
Query_10012_PhaC_Class_III__Haloferax_mediterranei_(AN:WP_004060157.1)
Query_10013_PhaC_Class_III__Haloquadratum_walsbyi_(AN:CCC40396.1)
Query_10014_PhaC_Class_III__Synechocystis_sp._PCC_6803_(AN:BAM51158.1)
Query_10015_PhaC_Class_IV__Bacillus_cereus_(AN:AAW84266.2)
Query_10016_PhaC_Class_IV__Bacillus_megaterium_(AN:AAD05260.1)
70
80
90
Consensus
V F X G X X X L X L X E A L X Q N I A R X A A E N T G A L N A A

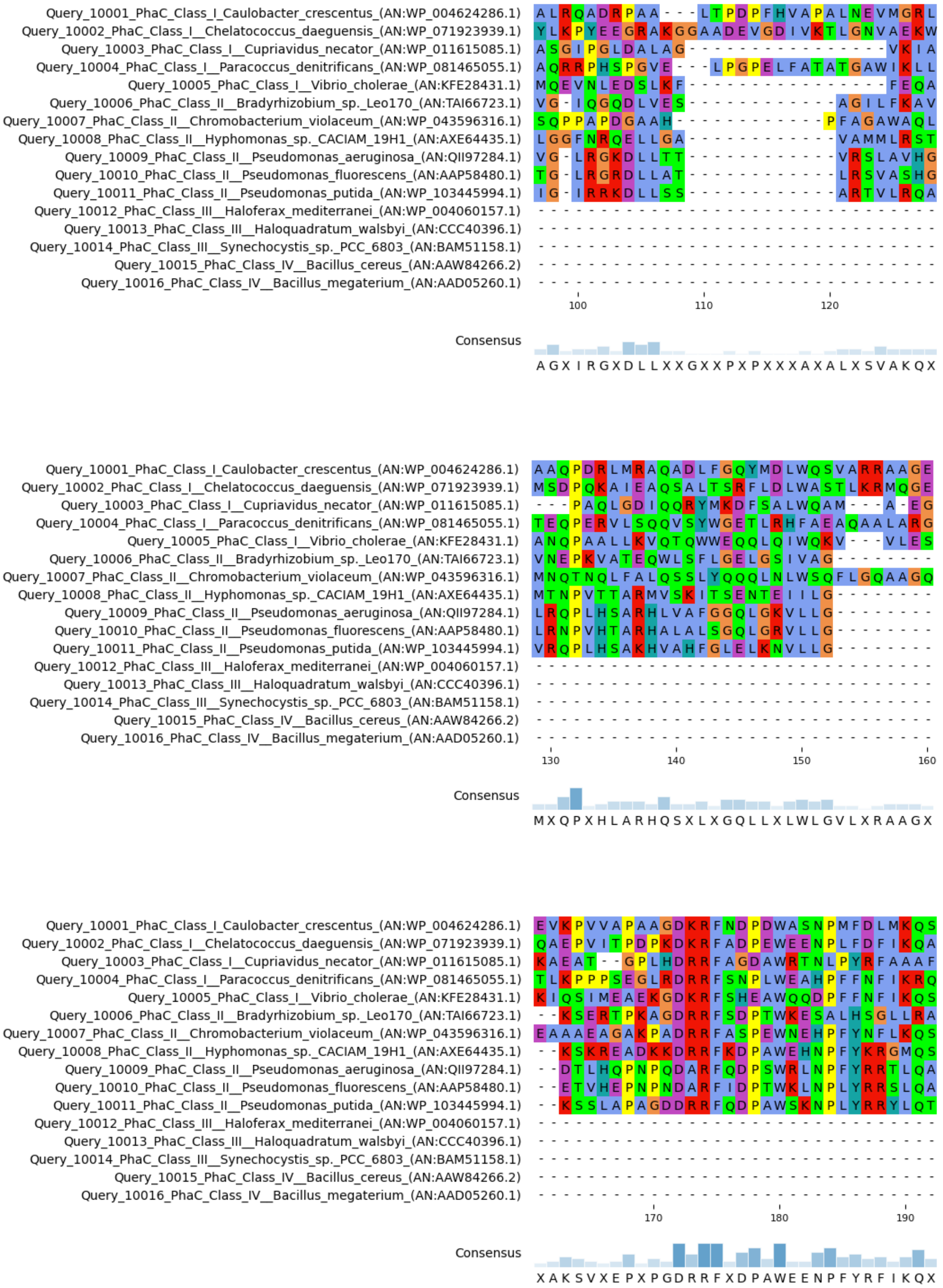
Query_10001_PhaC_Class_I_Caulobacter_crescentus_(AN:WP_004624286.1)
Query_10002_PhaC_Class_I__Chelatococcus_daeguensis_(AN:WP_071923939.1)
Query_10003_PhaC_Class_I__Cupriavidus_necator_(AN:WP_011615085.1)
Query_10004_PhaC_Class_I__Paracoccus_denitrificans_(AN:WP_081465055.1)
Query_10005_PhaC_Class_I__Vibrio_cholerae_(AN:KFE28431.1)
Query_10006_PhaC_Class_II__Bradyrhizobium_sp._Leo170_(AN:TAI66723.1)
Query_10007_PhaC_Class_II__Chromobacterium_violaceum_(AN:WP_043596316.1)
Query_10008_PhaC_Class_II__Hyphomonas_sp._CACIAM_19H1_(AN:AXE64435.1)
Query_10009_PhaC_Class_II__Pseudomonas_aeruginosa_(AN:QII97284.1)
Query_10010_PhaC_Class_II__Pseudomonas_fluorescens_(AN:AAP58480.1)
Query_10011_PhaC_Class_II__Pseudomonas_putida_(AN:WP_103445994.1)
Query_10012_PhaC_Class_III__Haloferax_mediterranei_(AN:WP_004060157.1)
Query_10013_PhaC_Class_III__Haloquadratum_walsbyi_(AN:CCC40396.1)
Query_10014_PhaC_Class_III__Synechocystis_sp._PCC_6803_(AN:BAM51158.1)
Query_10015_PhaC_Class_IV__Bacillus_cereus_(AN:AAW84266.2)
Query_10016_PhaC_Class_IV__Bacillus_megaterium_(AN:AAD05260.1)
100
110
120
Consensus
A G X I R G X D L L X X G X X P X P X X X A X A L X S V A K Q X
130
140
150
160
Consensus
M X Q P X H L A R H Q S X L X G Q L L X L W L G V L X R A A G X
170
180
190
Consensus
X A K S V X E P X P G D R R F X D P A W E E N P F Y R F I K Q X

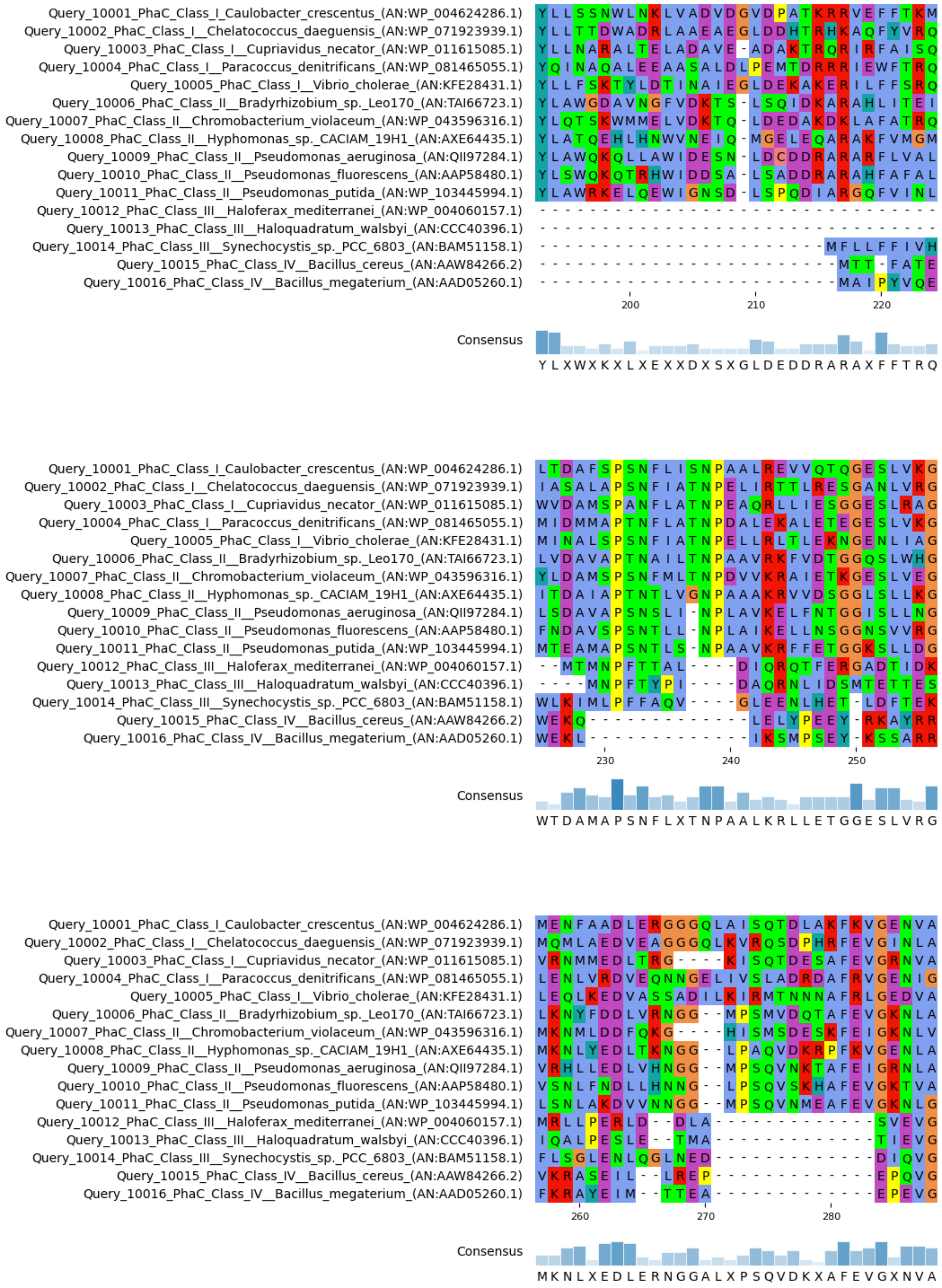
Query_10001_PhaC_Class_I_Caulobacter_crescentus_(AN:WP_004624286.1)
Query_10002_PhaC_Class_I__Chelatococcus_daeguensis_(AN:WP_071923939.1)
Query_10003_PhaC_Class_I__Cupriavidus_necator_(AN:WP_011615085.1)
Query_10004_PhaC_Class_I__Paracoccus_denitrificans_(AN:WP_081465055.1)
Query_10005_PhaC_Class_I__Vibrio_cholerae_(AN:KFE28431.1)
Query_10006_PhaC_Class_II__Bradyrhizobium_sp._Leo170_(AN:TAI66723.1)
Query_10007_PhaC_Class_II__Chromobacterium_violaceum_(AN:WP_043596316.1)
Query_10008_PhaC_Class_II__Hyphomonas_sp._CACIAM_19H1_(AN:AXE64435.1)
Query_10009_PhaC_Class_II__Pseudomonas_aeruginosa_(AN:QII97284.1)
Query_10010_PhaC_Class_II__Pseudomonas_fluorescens_(AN:AAP58480.1)
Query_10011_PhaC_Class_II__Pseudomonas_putida_(AN:WP_103445994.1)
Query_10012_PhaC_Class_III__Haloferax_mediterranei_(AN:WP_004060157.1)
Query_10013_PhaC_Class_III__Haloquadratum_walsbyi_(AN:CCC40396.1)
Query_10014_PhaC_Class_III__Synechocystis_sp._PCC_6803_(AN:BAM51158.1)
Query_10015_PhaC_Class_IV__Bacillus_cereus_(AN:AAW84266.2)
Query_10016_PhaC_Class_IV__Bacillus_megaterium_(AN:AAD05260.1)
200
210
220
Consensus
Y L X W X K X L X E X X D X S X G L D E D D R A R A X F F T R Q
230
240
250
Consensus
W T D A M A P S N F L X T N P A A L K R L L E T G G E S L V R G
260
270
280
Consensus
M K N L X E D L E R N G G A L X P S Q V D K X A F E V G X N V A

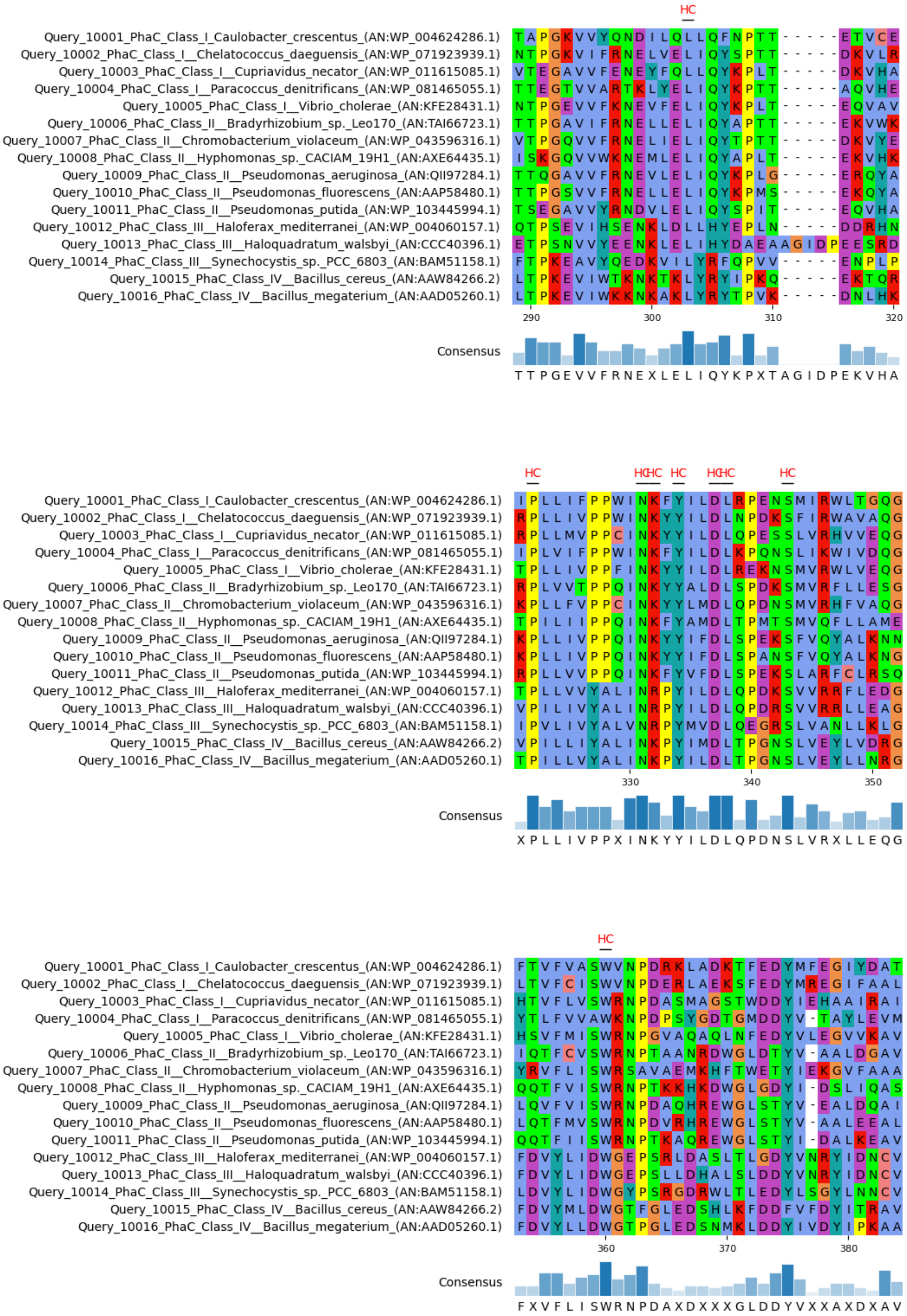
HC
Query_10001_PhaC_Class_I_Caulobacter_crescentus_(AN:WP_004624286.1)
Query_10002_PhaC_Class_I__Chelatococcus_daeguensis_(AN:WP_071923939.1)
Query_10003_PhaC_Class_I__Cupriavidus_necator_(AN:WP_011615085.1)
Query_10004_PhaC_Class_I__Paracoccus_denitrificans_(AN:WP_081465055.1)
Query_10005_PhaC_Class_I__Vibrio_cholerae_(AN:KFE28431.1)
Query_10006_PhaC_Class_II__Bradyrhizobium_sp._Leo170_(AN:TAI66723.1)
Query_10007_PhaC_Class_II__Chromobacterium_violaceum_(AN:WP_043596316.1)
Query_10008_PhaC_Class_II__Hyphomonas_sp._CACIAM_19H1_(AN:AXE64435.1)
Query_10009_PhaC_Class_II__Pseudomonas_aeruginosa_(AN:QII97284.1)
Query_10010_PhaC_Class_II__Pseudomonas_fluorescens_(AN:AAP58480.1)
Query_10011_PhaC_Class_II__Pseudomonas_putida_(AN:WP_103445994.1)
Query_10012_PhaC_Class_III__Haloferax_mediterranei_(AN:WP_004060157.1)
Query_10013_PhaC_Class_III__Haloquadratum_walsbyi_(AN:CCC40396.1)
Query_10014_PhaC_Class_III__Synechocystis_sp._PCC_6803_(AN:BAM51158.1)
Query_10015_PhaC_Class_IV__Bacillus_cereus_(AN:AAW84266.2)
Query_10016_PhaC_Class_IV__Bacillus_megaterium_(AN:AAD05260.1)
290 300 310 320
Consensus
T T P G E V V F R N E X L E L I Q Y K P X T A G I D P E K V H A
HC HCHC HC HCHC HC
330 340 350
Consensus
X P L L I V P P X I N K Y Y I L D L Q P D N S L V R X L L E Q G
HC
360 370 380
Consensus
F X V F L I S W R N P D A X D X X X G L D D Y V X X A X D X A V

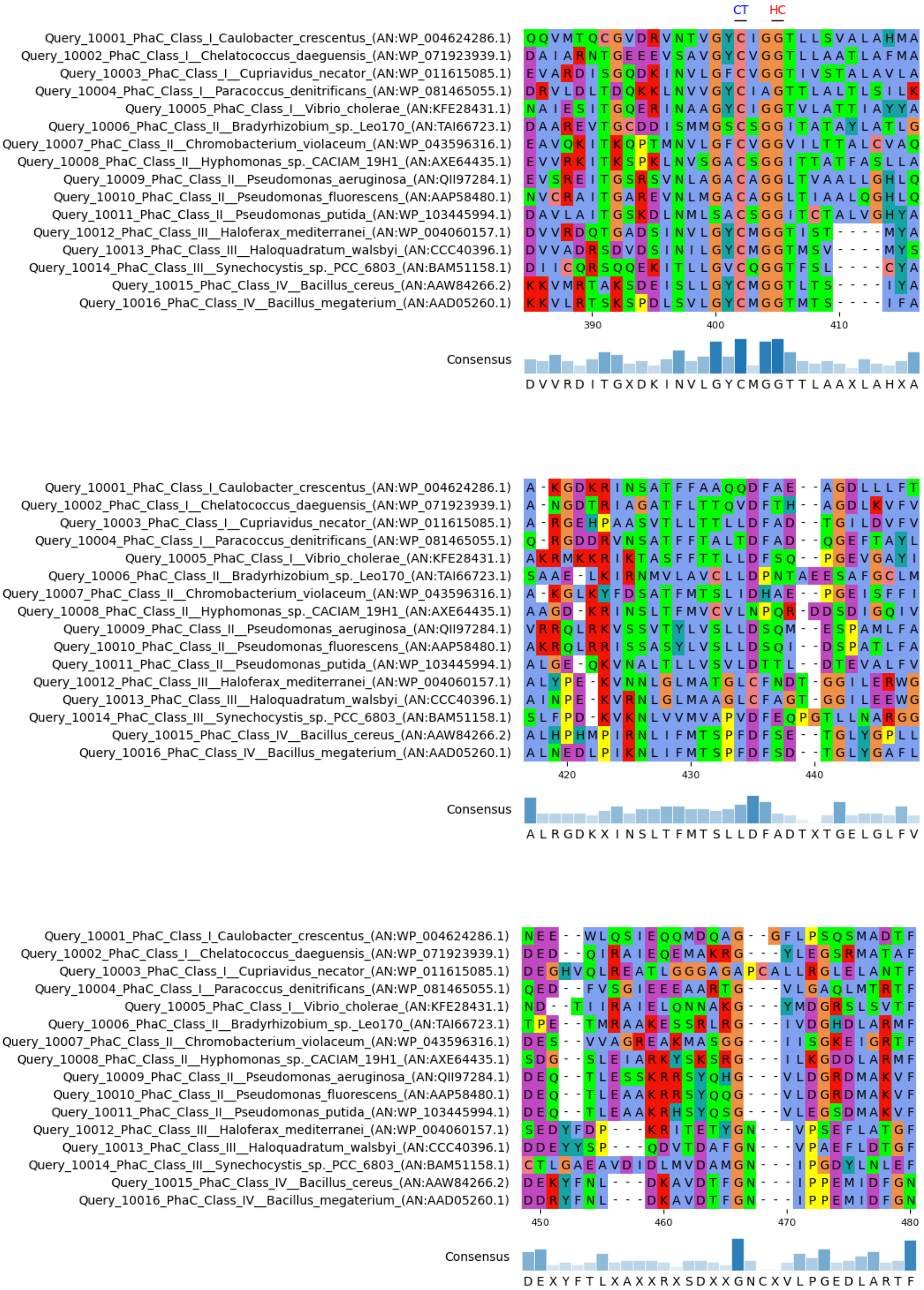
CT
HC
Query_10001_PhaC_Class_I_Caulobacter_crescentus_(AN:WP_004624286.1)
Query_10002_PhaC_Class_I__Chelatococcus_daeguensis_(AN:WP_071923939.1)
Query_10003_PhaC_Class_I__Cupriavidus_necator_(AN:WP_011615085.1)
Query_10004_PhaC_Class_I__Paracoccus_denitrificans_(AN:WP_081465055.1)
Query_10005_PhaC_Class_I__Vibrio_cholerae_(AN:KFE28431.1)
Query_10006_PhaC_Class_II__Bradyrhizobium_sp._Leo170_(AN:TAI66723.1)
Query_10007_PhaC_Class_II__Chromobacterium_violaceum_(AN:WP_043596316.1)
Query_10008_PhaC_Class_II__Hyphomonas_sp._CACIAM_19H1_(AN:AXE64435.1)
Query_10009_PhaC_Class_II__Pseudomonas_aeruginosa_(AN:QII97284.1)
Query_10010_PhaC_Class_II__Pseudomonas_fluorescens_(AN:AAP58480.1)
Query_10011_PhaC_Class_II__Pseudomonas_putida_(AN:WP_103445994.1)
Query_10012_PhaC_Class_III__Haloferax_mediterranei_(AN:WP_004060157.1)
Query_10013_PhaC_Class_III__Haloquadratum_walsbyi_(AN:CCC40396.1)
Query_10014_PhaC_Class_III__Synechocystis_sp._PCC_6803_(AN:BAM51158.1)
Query_10015_PhaC_Class_IV__Bacillus_cereus_(AN:AAW84266.2)
Query_10016_PhaC_Class_IV__Bacillus_megaterium_(AN:AAD05260.1)
390
400
410
Consensus
D V V R D I T G X D K I N V L G Y C M G G T T L A A X L A H X A
420
430
440
Consensus
A L R G D K X I N S L T F M T S L L D F A D T X T G E L G L F V
450
460
470
480
Consensus
D E X Y F T L X A X X R X S D X X G N C X V L P G E D L A R T F

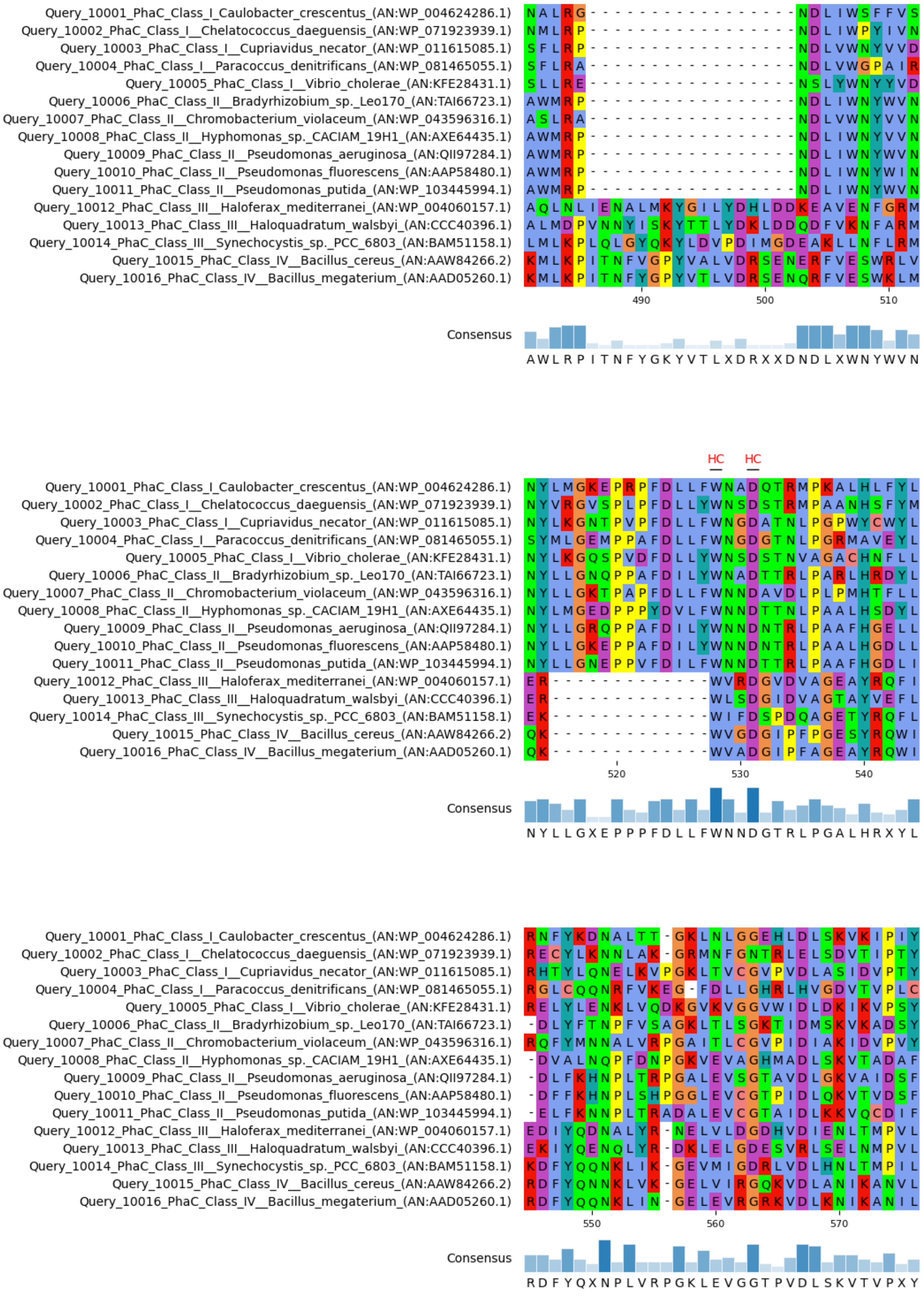
Query_10001_PhaC_Class_I_Caulobacter_crescentus_(AN:WP_004624286.1)
Query_10002_PhaC_Class_I__Chelatococcus_daeguensis_(AN:WP_071923939.1)
Query_10003_PhaC_Class_I__Cupriavidus_necator_(AN:WP_011615085.1)
Query_10004_PhaC_Class_I__Paracoccus_denitrificans_(AN:WP_081465055.1)
Query_10005_PhaC_Class_I__Vibrio_cholerae_(AN:KFE28431.1)
Query_10006_PhaC_Class_II__Bradyrhizobium_sp._Leo170_(AN:TAI66723.1)
Query_10007_PhaC_Class_II__Chromobacterium_violaceum_(AN:WP_043596316.1)
Query_10008_PhaC_Class_II__Hyphomonas_sp._CACIAM_19H1_(AN:AXE64435.1)
Query_10009_PhaC_Class_II__Pseudomonas_aeruginosa_(AN:QII97284.1)
Query_10010_PhaC_Class_II__Pseudomonas_fluorescens_(AN:AAP58480.1)
Query_10011_PhaC_Class_II__Pseudomonas_putida_(AN:WP_103445994.1)
Query_10012_PhaC_Class_III__Haloferax_mediterranei_(AN:WP_004060157.1)
Query_10013_PhaC_Class_III__Haloquadratum_walsbyi_(AN:CCC40396.1)
Query_10014_PhaC_Class_III__Synechocystis_sp._PCC_6803_(AN:BAM51158.1)
Query_10015_PhaC_Class_IV__Bacillus_cereus_(AN:AAW84266.2)
Query_10016_PhaC_Class_IV__Bacillus_megaterium_(AN:AAD05260.1)
490
500
510
Consensus
A W L R P I T N F Y G K Y V T L X D R X X D N D L X W N Y W V N
HC
HC
520
530
540
Consensus
N Y L L G X E P P P F D L L F W N N D G T R L P G A L H R X Y L
550
560
570
Consensus
R D F Y Q X N P L V R P G K L E V G G T P V D L S K V T V P X Y

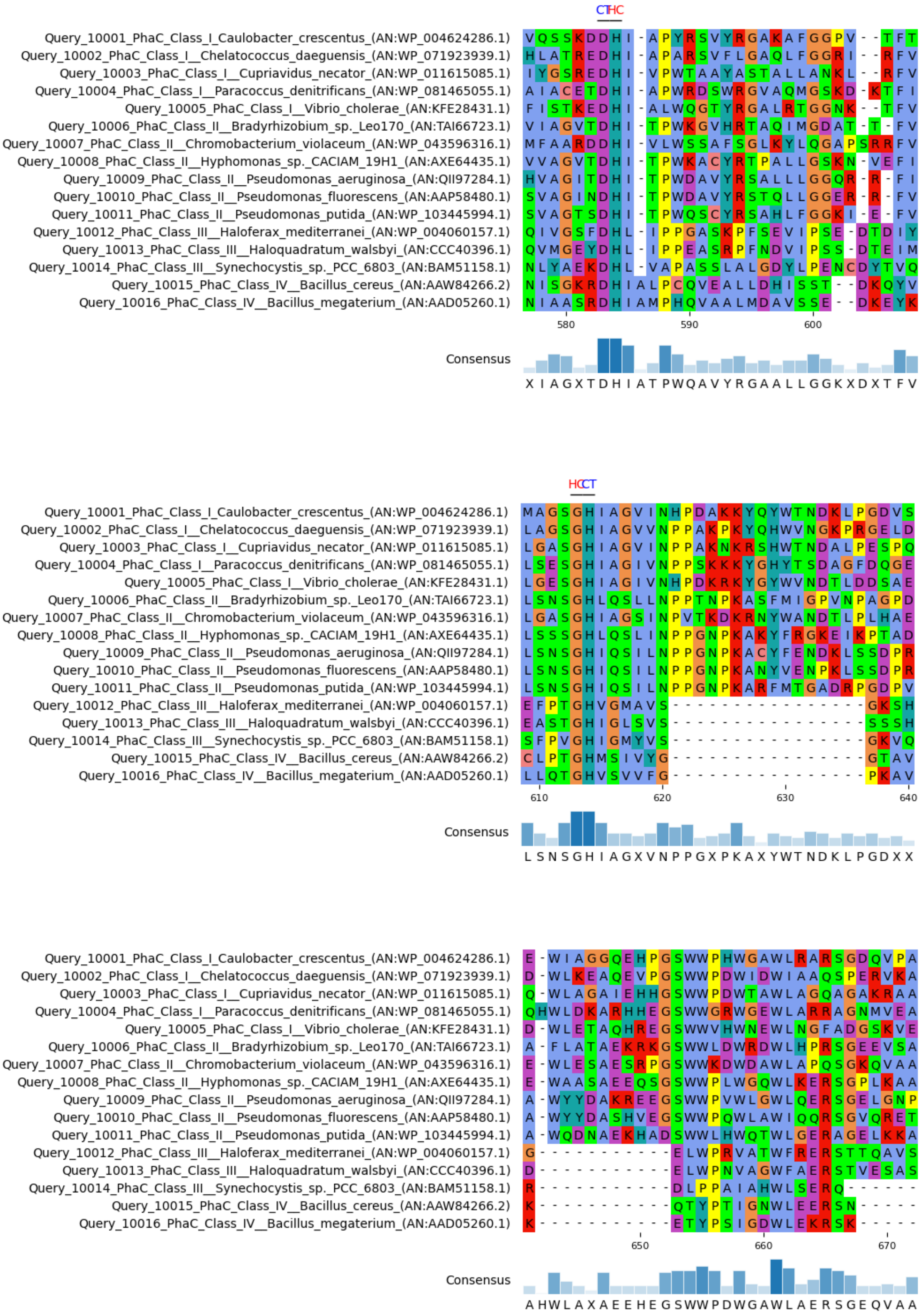

CTHC
Query_10001_PhaC_Class_I_Caulobacter_crescentus_(AN:WP_004624286.1)
Query_10002_PhaC_Class_I__Chelatococcus_daeguensis_(AN:WP_071923939.1)
Query_10003_PhaC_Class_I__Cupriavidus_necator_(AN:WP_011615085.1)
Query_10004_PhaC_Class_I__Paracoccus_denitrificans_(AN:WP_081465055.1)
Query_10005_PhaC_Class_I__Vibrio_cholerae_(AN:KFE28431.1)
Query_10006_PhaC_Class_II__Bradyrhizobium_sp._Leo170_(AN:TAI66723.1)
Query_10007_PhaC_Class_II__Chromobacterium_violaceum_(AN:WP_043596316.1)
Query_10008_PhaC_Class_II__Hyphomonas_sp._CACIAM_19H1_(AN:AXE64435.1)
Query_10009_PhaC_Class_II__Pseudomonas_aeruginosa_(AN:QII97284.1)
Query_10010_PhaC_Class_II__Pseudomonas_fluorescens_(AN:AAP58480.1)
Query_10011_PhaC_Class_II__Pseudomonas_putida_(AN:WP_103445994.1)
Query_10012_PhaC_Class_III__Haloferax_mediterranei_(AN:WP_004060157.1)
Query_10013_PhaC_Class_III__Haloquadratum_walsbyi_(AN:CCC40396.1)
Query_10014_PhaC_Class_III__Synechocystis_sp._PCC_6803_(AN:BAM51158.1)
Query_10015_PhaC_Class_IV__Bacillus_cereus_(AN:AAW84266.2)
Query_10016_PhaC_Class_IV__Bacillus_megaterium_(AN:AAD05260.1)
580
590
600
Consensus
X I A G X T D H I A T P W Q A V Y R G A A L L G G K X D X T F V
HCCT
610
620
630
640
Consensus
L S N S G H I A G X V N P P G X P K A X Y W T N D K L P G D X X
650
660
670
Consensus
A H W L A X A E E H E G S W W P D W G A W L A E R S G E Q V A A

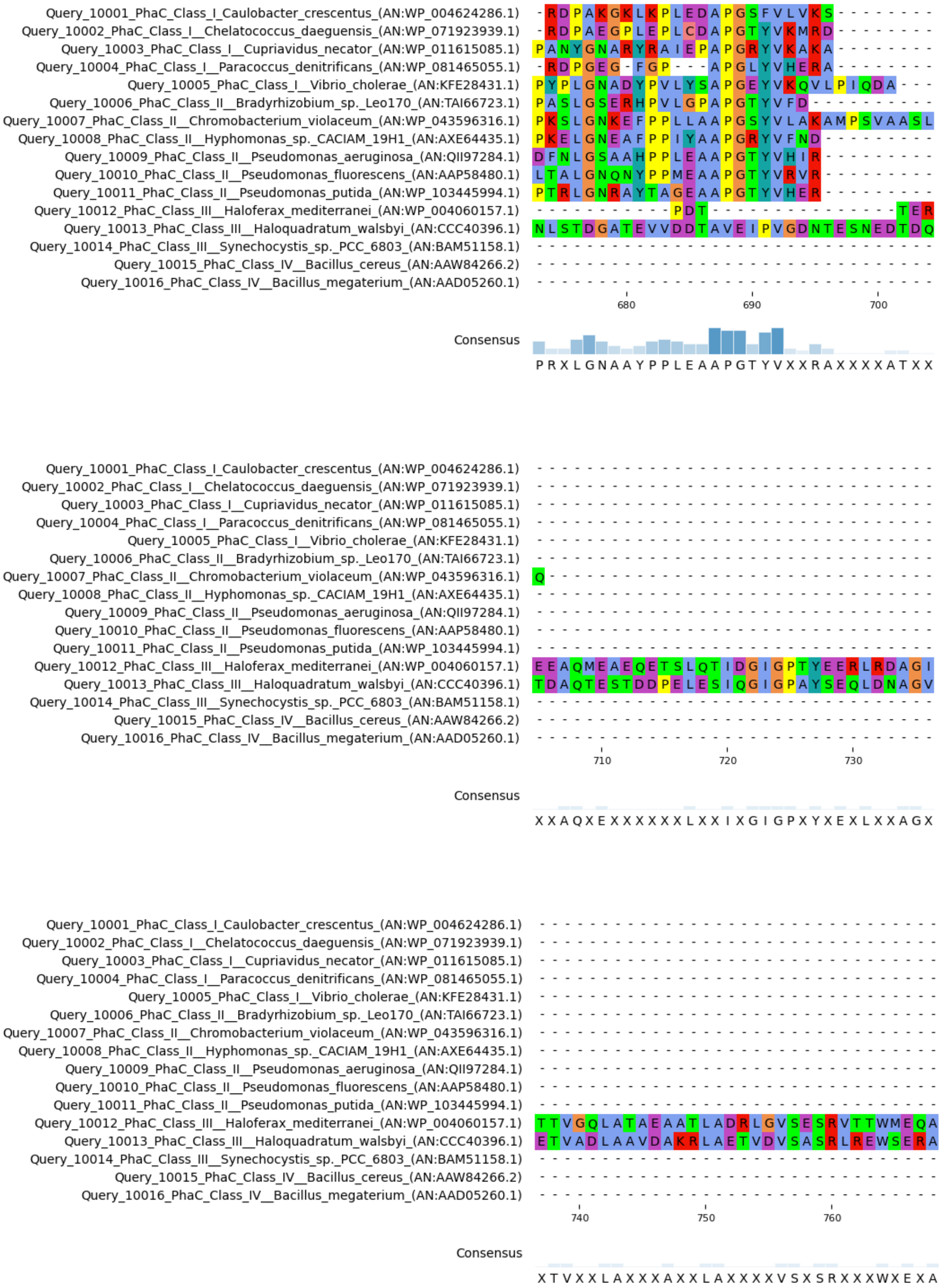
Query_10001_PhaC_Class_I_Caulobacter_crescentus_(AN:WP_004624286.1)
Query_10002_PhaC_Class_I__Chelatococcus_daeguensis_(AN:WP_071923939.1)
Query_10003_PhaC_Class_I__Cupriavidus_necator_(AN:WP_011615085.1)
Query_10004_PhaC_Class_I__Paracoccus_denitrificans_(AN:WP_081465055.1)
Query_10005_PhaC_Class_I__Vibrio_cholerae_(AN:KFE28431.1)
Query_10006_PhaC_Class_II__Bradyrhizobium_sp._Leo170_(AN:TAI66723.1)
Query_10007_PhaC_Class_II__Chromobacterium_violaceum_(AN:WP_043596316.1)
Query_10008_PhaC_Class_II__Hyphomonas_sp._CACIAM_19H1_(AN:AXE64435.1)
Query_10009_PhaC_Class_II__Pseudomonas_aeruginosa_(AN:QII97284.1)
Query_10010_PhaC_Class_II__Pseudomonas_fluorescens_(AN:AAP58480.1)
Query_10011_PhaC_Class_II__Pseudomonas_putida_(AN:WP_103445994.1)
Query_10012_PhaC_Class_III__Haloferax_mediterranei_(AN:WP_004060157.1)
Query_10013_PhaC_Class_III__Haloquadratum_walsbyi_(AN:CCC40396.1)
Query_10014_PhaC_Class_III__Synechocystis_sp._PCC_6803_(AN:BAM51158.1)
Query_10015_PhaC_Class_IV__Bacillus_cereus_(AN:AAW84266.2)
Query_10016_PhaC_Class_IV__Bacillus_megaterium_(AN:AAD05260.1)
680
690
700
Consensus
P R X L G N A A Y P P L E A A P G T Y V X X R A X X X X X A T X X
710
720
730
Consensus
X X A Q X E X X X X X X L X X I X G I G P X Y X E X L X X A G X
740
750
760
Consensus
X T V X X L A X X X A X X L A X X X X X V S X S R X X X W X E X A

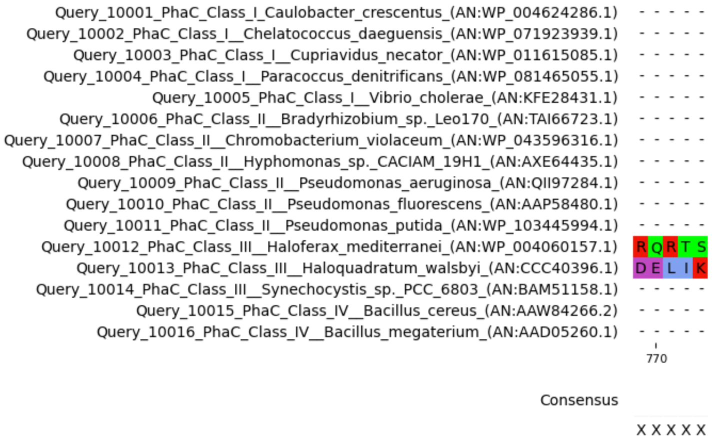


**Supplementary Figure S1**: Multiple sequence alignment of the 16 well-described or automatically annotated PhaC sequences. MSA was performed with COBALT and visualized with PYMSAVIZ. Hyperconserved residues are indicated with a red HC on top of the column. Residues part of the catalytic triad are indicated with a blue CT on top of the column.

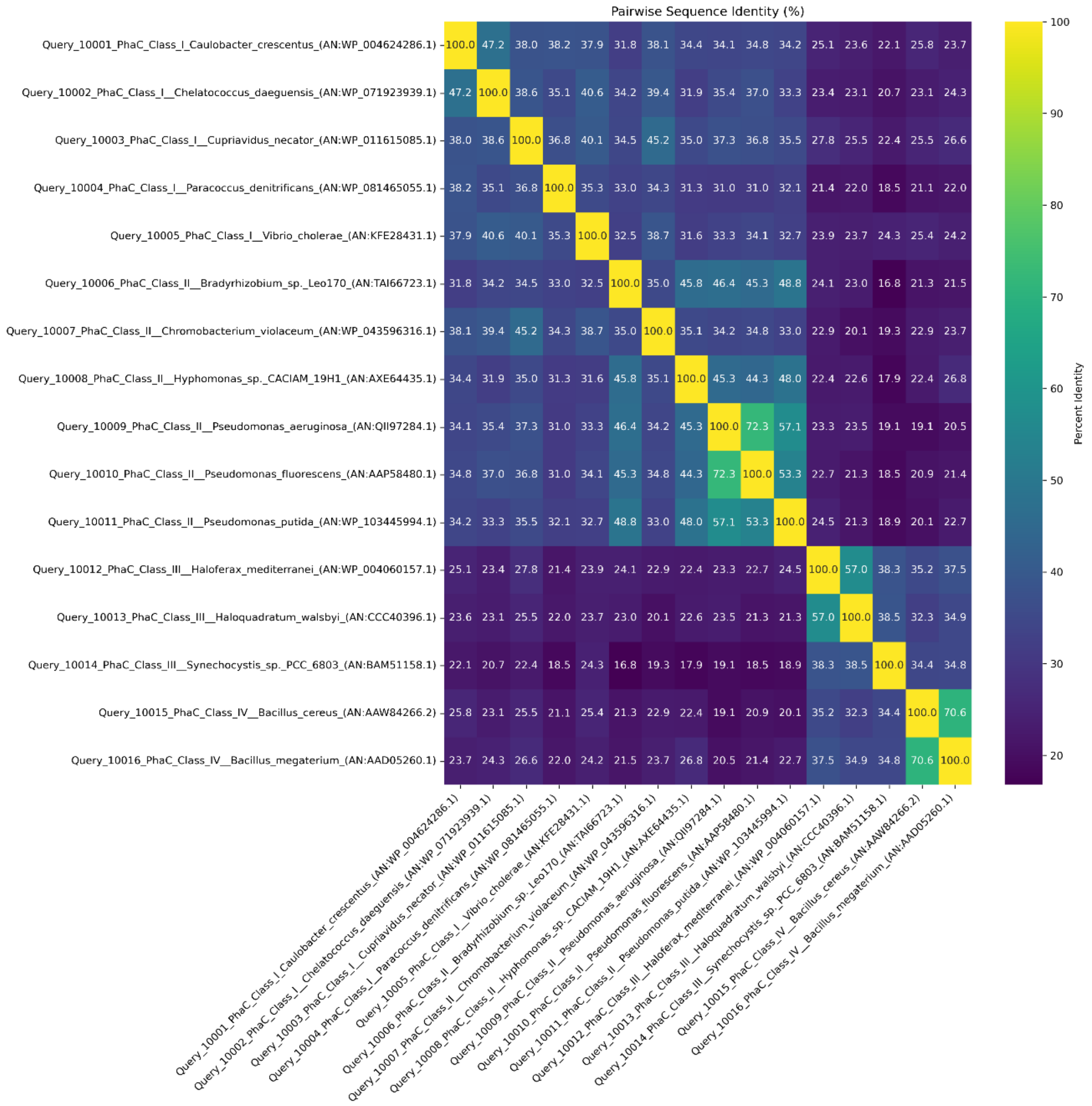


**Supplementary figure S2**: Pairwise percent identity matrix of aligned PhaC protein sequences. Percent identity was calculated based on non-gap aligned positions for each sequence pair. The color scale represents sequence similarity, with lighter shades indicating higher identity. The analysis was performed using Python (Biopython, Seaborn, Matplotlib), based on a multiple sequence alignment generated with COBALT.

**Supplementary Table S2**: Filtered list of putative PhaC sequences in bacterial moderate thermophiles. This list is filtered on the lipase-like box location and the conservation of the hyperconserved residues.

| Accession | Annotation | Microorganism |
|---|---|---|
| **WP_132806208.1** | class I poly(R)-hydroxyalkanoic acid synthase | *Tepidamorphus gemmatus* |
| **WP_028875124.1** | class I poly(R)-hydroxyalkanoic acid synthase | *Tepidiphilus margaritifer* |
| **WP_142804171.1** | class I poly(R)-hydroxyalkanoic acid synthase | *Tepidiphilus sp. J10* |
| **WP_141056142.1** | class I poly(R)-hydroxyalkanoic acid synthase | *Tepidiphilus succinatimandens* |
| **WP_055423166.1** | class I poly(R)-hydroxyalkanoic acid synthase | *Tepidiphilus thermophilus* |

| | | |
|---|---|---|
| **WP_169116172.1** | class I poly(R)-hydroxyalkanoic acid synthase | *Tepidiphilus baoligensis* |
| **WP_221928882.1** | class I poly(R)-hydroxyalkanoic acid synthase | *Tepidiphilus olei* |
| **WP_119062396.1** | class I poly(R)-hydroxyalkanoic acid synthase | *Dichotomicrobium thermohalophilum* |
| **WP_119334366.1** | class I poly(R)-hydroxyalkanoic acid synthase | *Hydrogenophilus thermoluteolus* |
| **WP_217126752.1** | class I poly(R)-hydroxyalkanoic acid synthase | *Hydrogenophilus thiooxidans* |
| **WP_219973294.1** | class I poly(R)-hydroxyalkanoic acid synthase | *Hydrogenophilus thermoluteolus* |
| **WP_206202189.1** | class I poly(R)-hydroxyalkanoic acid synthase | *Tepidiphilus succinatimandens* |
| **WP_051240655.1** | class I poly(R)-hydroxyalkanoic acid synthase | *Tepidiphilus margaritifer* |
| **WP_055423697.1** | class I poly(R)-hydroxyalkanoic acid synthase | *Tepidiphilus thermophilus* |
| **NMH16349.1** | class I poly(R)-hydroxyalkanoic acid synthase | *Tepidiphilus baoligensis* |
| **WP_178086736.1** | class I poly(R)-hydroxyalkanoic acid synthase | *Tepidiphilus baoligensis* |
| **WP_240635972.1** | class I poly(R)-hydroxyalkanoic acid synthase | *Caldimonas tepidiphila* |
| **WP_142808715.1** | class I poly(R)-hydroxyalkanoic acid synthase | *Tepidiphilus olei* |
| **WP_238947598.1** | class I poly(R)-hydroxyalkanoic acid synthase | *Caldimonas brevitalea* |
| **WP_142803501.1** | class I poly(R)-hydroxyalkanoic acid synthase | *Tepidiphilus sp. J10* |
| **WP_245876440.1** | class I poly(R)-hydroxyalkanoic acid synthase | *Caldimonas caldifontis* |
| **PPE66523.1** | class I poly(R)-hydroxyalkanoic acid synthase | *Caldimonas caldifontis* |
| **WP_232300999.1** | class I poly(R)-hydroxyalkanoic acid synthase | *Caldimonas taiwanensis* |
| **WP_026330149.1** | class I poly(R)-hydroxyalkanoic acid synthase | *Caldimonas manganoxidans* |
| **WP_264894475.1** | class I poly(R)-hydroxyalkanoic acid synthase | *Schlegelella aquatica* |
| **WP_232529522.1** | class I poly(R)-hydroxyalkanoic acid synthase | *Caldimonas thermodepolymerans* |
| **WP_251780501.1** | class I poly(R)-hydroxyalkanoic acid synthase | *Schlegelella sp. S2-27* |
| **MBA3794682.1** | class I poly(R)-hydroxyalkanoic acid synthase | *Rubrobacter sp.* |
| **PPE70723.1** | class I poly(R)-hydroxyalkanoic acid synthase | *Caldimonas thermodepolymerans* |
| **QIN85027.1** | class I poly(R)-hydroxyalkanoic acid synthase | *Rubrobacter tropicus* |
| **WP_219971775.1** | class I poly(R)-hydroxyalkanoic acid synthase | *Hydrogenophilus thermoluteolus* |
| **WP_228282310.1** | class I poly(R)-hydroxyalkanoic acid synthase | *Rubrobacter tropicus* |
| **WP_197713626.1** | class I poly(R)-hydroxyalkanoic acid synthase | *Hydrogenophilus thermoluteolus* |
| **WP_144324707.1** | class I poly(R)-hydroxyalkanoic acid synthase | *Tepidimonas aquatica* |
| **WP_132961516.1** | class I poly(R)-hydroxyalkanoic acid synthase | *Tepidimonas ignava* |
| **WP_058616235.1** | class I poly(R)-hydroxyalkanoic acid synthase | *Tepidimonas taiwanensis* |
| **WP_143892980.1** | class I poly(R)-hydroxyalkanoic acid synthase | *Tepidimonas sediminis* |
| **WP_217125565.1** | class I poly(R)-hydroxyalkanoic acid synthase | *Hydrogenophilus thiooxidans* |
| **WP_176067442.1** | class I poly(R)-hydroxyalkanoic acid synthase | *Schlegelella koreensis* |

| | | |
|---|---|---|
| **WP_143902344.1** | class I poly(R)-hydroxyalkanoic acid synthase | *Tepidimonas thermarum* |
| **WP_224441037.1** | class I poly(R)-hydroxyalkanoic acid synthase | *Tepidimonas taiwanensis* |
| **MBT0745838.1** | class I poly(R)-hydroxyalkanoic acid synthase | *Anaerolineae bacterium* |
| **WP_231572220.1** | class I poly(R)-hydroxyalkanoic acid synthase | *Tepidimonas taiwanensis* |
| **WP_143968067.1** | class I poly(R)-hydroxyalkanoic acid synthase | *Tepidimonas fonticaldi* |
| **GIX25162.1** | MAG: poly-beta-hydroxybutyrate polymerase | *Caldimonas sp.* |
| **WP_082955490.1** | class I poly(R)-hydroxyalkanoic acid synthase | *Tepidimonas fonticaldi* |
| **MBA2692584.1** | class I poly(R)-hydroxyalkanoic acid synthase | *Rubrobacter sp.* |
| **WP_144327474.1** | class I poly(R)-hydroxyalkanoic acid synthase | *Tepidimonas charontis* |
| **MBA2441635.1** | class I poly(R)-hydroxyalkanoic acid synthase | *Rubrobacter sp.* |
| **MBL6982746.1** | class I poly(R)-hydroxyalkanoic acid synthase | *Anaerolineales bacterium* |
| **WP_132688328.1** | class I poly(R)-hydroxyalkanoic acid synthase | *Rubrobacter taiwanensis* |
| **WP_219975642.1** | class I poly(R)-hydroxyalkanoic acid synthase | *Rubrobacter xylanophilus* |
| **WP_276706894.1** | class I poly(R)-hydroxyalkanoic acid synthase | *Thiomonas arsenitoxydans* |
| **WP_038682553.1** | class I poly(R)-hydroxyalkanoic acid synthase | *Rubrobacter radiotolerans* |
| **WP_276729874.1** | class I poly(R)-hydroxyalkanoic acid synthase | *Thiomonas arsenitoxydans* |
| **OZB71695.1** | MAG: class I poly(R)-hydroxyalkanoic acid synthase | *Thiomonas sp. 13-64-67* |
| **MBA2376518.1** | class I poly(R)-hydroxyalkanoic acid synthase | *Rubrobacter sp.* |
| **WP_156052721.1** | MULTISPECIES: class I poly(R)-hydroxyalkanoic acid synthase | *unclassified Thiomonas* |
| **WP_041608940.1** | MULTISPECIES: class I poly(R)-hydroxyalkanoic acid synthase | *Thiomonas* |
| **MCA3748477.1** | class I poly(R)-hydroxyalkanoic acid synthase | *Rubrobacter sp.* |
| **WP_055450956.1** | class I poly(R)-hydroxyalkanoic acid synthase | *Thiomonas bhubaneswarensis* |
| **WP_240432455.1** | class I poly(R)-hydroxyalkanoic acid synthase | *Rubrobacter indicoceani* |
| **WP_246100573.1** | class I poly(R)-hydroxyalkanoic acid synthase | *Tepidimonas alkaliphilus* |
| **CUA98300.1** | poly(R)-hydroxyalkanoic acid synthase, class I | *Thiomonas bhubaneswarensis* |
| **MCB9135560.1** | class I poly(R)-hydroxyalkanoic acid synthase | *Anaerolineales bacterium* |
| **ODU97219.1** | MAG: class I poly(R)-hydroxyalkanoic acid synthase | *Thiomonas sp. SCN 64-16* |
| **CDW95509.1** | Poly-beta-hydroxybutyrate polymerase | *Thiomonas sp. CB2* |
| **CAZ88317.1** | Poly-beta-hydroxybutyrate polymerase (Poly(3-hydroxybutyrate) polymerase) (PHB polymerase) (PHB synthase) (Poly(3-hydroxyalkanoate) polymerase) (PHA polymerase) (PHA synthase) (Polyhydroxyalkanoic acid synthase) | *Thiomonas arsenitoxydans* |
| **WP_143527770.1** | class I poly(R)-hydroxyalkanoic acid synthase | *Rubrobacter xylanophilus* |
| **WP_011565327.1** | class I poly(R)-hydroxyalkanoic acid synthase | *Rubrobacter xylanophilus* |
| **NUM46192.1** | class I poly(R)-hydroxyalkanoic acid synthase | *Anaerolineales bacterium* |
| **OYV30990.1** | MAG: class I poly(R)-hydroxyalkanoic acid synthase | *Thiomonas sp. 20-64-9* |

| | | |
|---|---|---|
| **OZB45668.1** | MAG: class I poly(R)-hydroxyalkanoic acid synthase | *Thiomonas sp. 15-66-11* |
| **WP_094160349.1** | class I poly(R)-hydroxyalkanoic acid synthase | *Thiomonas delicata* |
| **WP_112485572.1** | alpha/beta fold hydrolase | *Thiomonas sp. X19* |
| **MBA2691072.1** | class I poly(R)-hydroxyalkanoic acid synthase | *Rubrobacter sp.* |
| **OZB63021.1** | MAG: class I poly(R)-hydroxyalkanoic acid synthase | *Thiomonas sp. 13-66-29* |
| **WP_079419847.1** | class I poly(R)-hydroxyalkanoic acid synthase | *Thiomonas intermedia* |
| **WP_038168446.1** | alpha/beta fold hydrolase | *Thiomonas sp. FB-Cd* |
| **MCE1161470.1** | class I poly(R)-hydroxyalkanoic acid synthase | *Thiomonas sp.* |
| **WP_072756679.1** | alpha/beta fold hydrolase | *Thermomonas hydrothermalis* |
| **WP_132766364.1** | alpha/beta fold hydrolase | *Caldimonas thermodepolymerans* |
| **WP_072030302.1** | alpha/beta fold hydrolase | *Amycolatopsis keratiniphila* |
| **WP_013228250.1** | alpha/beta fold hydrolase | *Amycolatopsis mediterranei* |
| **NMH15642.1** | alpha/beta fold hydrolase | *Tepidiphilus baoligensis* |
| **WP_132805903.1** | alpha/beta fold hydrolase | *Tepidamorphus gemmatus* |
| **WP_221928946.1** | MULTISPECIES: alpha/beta fold hydrolase | *Tepidiphilus* |
| **WP_178086720.1** | alpha/beta fold hydrolase | *Tepidiphilus baoligensis* |
| **WP_038166483.1** | alpha/beta fold hydrolase | *Thiomonas sp. FB-6* |
| **WP_021249554.1** | MULTISPECIES: alpha/beta fold hydrolase | *Pseudomonadota* |
| **WP_211218511.1** | alpha/beta fold hydrolase | *Tepidiphilus margaritifer* |
| **WP_033289844.1** | alpha/beta fold hydrolase | *Amycolatopsis jejuensis* |
| **WP_141055074.1** | alpha/beta fold hydrolase | *Tepidiphilus succinatimandens* |
| **WP_198289022.1** | alpha/beta fold hydrolase | *Tepidiphilus thermophilus* |
| **MBP6998508.1** | alpha/beta fold hydrolase | *Tepidiphilus sp.* |
| **WP_276706597.1** | alpha/beta fold hydrolase | *Thiomonas arsenitoxydans* |
| **WP_276732369.1** | alpha/beta fold hydrolase | *Thiomonas arsenitoxydans* |
| **WP_013105489.1** | MULTISPECIES: alpha/beta fold hydrolase | *Thiomonas* |
| **OYV30273.1** | MAG: poly-beta-hydroxybutyrate polymerase | *Thiomonas sp. 20-64-9* |
| **WP_156055210.1** | MULTISPECIES: alpha/beta fold hydrolase | *unclassified Thiomonas* |
| **MBA3952794.1** | alpha/beta fold hydrolase | *Rubrobacter sp.* |
| **OZB77168.1** | MAG: poly-beta-hydroxybutyrate polymerase | *Thiomonas sp. 14-64-326* |
| **WP_082454307.1** | alpha/beta fold hydrolase | *Thiomonas bhubaneswarensis* |
| **WP_276706184.1** | alpha/beta fold hydrolase | *Thiomonas arsenitoxydans* |
| **CUA96569.1** | Poly(3-hydroxyalkanoate) synthetase | *Thiomonas bhubaneswarensis* |
| **MCS6810960.1** | class I poly(R)-hydroxyalkanoic acid synthase | *Tepidimonas sp.* |
| **WP_037359336.1** | alpha/beta fold hydrolase | *Amycolatopsis orientalis* |
| **CQR32534.1** | Poly-beta-hydroxybutyrate polymerase domain protein | *Thiomonas arsenitoxydans* |
| **NIV40667.1** | alpha/beta fold hydrolase | *Anaerolineae bacterium* |
| **WP_119060019.1** | alpha/beta fold hydrolase | *Dichotomicrobium thermohalophilum* |
| **MCE1163212.1** | alpha/beta fold hydrolase | *Thiomonas sp.* |
| **WP_079415230.1** | alpha/beta fold hydrolase | *Thiomonas intermedia* |
| **CDW95291.1** | Poly-beta-hydroxybutyrate polymerase domain protein | *Thiomonas sp. CB2* |
| **MBW7886120.1** | alpha/beta fold hydrolase | *Caldilineaceae bacterium* |
| **OZB51127.1** | MAG: poly-beta-hydroxybutyrate polymerase | *Thiomonas sp. 14-66-4* |
| **WP_081857951.1** | alpha/beta fold hydrolase | *Thiomonas sp. FB-Cd* |
| **WP_249419217.1** | alpha/beta fold hydrolase | *Hydrogenophilus thiooxidans* |
| **OZB64511.1** | MAG: poly-beta-hydroxybutyrate polymerase | *Thiomonas sp. 13-66-29* |
| **MBW7657116.1** | alpha/beta fold hydrolase | *Hydrogenophilus thermoluteolus* |
| **WP_197713757.1** | alpha/beta fold hydrolase | *Hydrogenophilus thermoluteolus* |
| **WP_094162151.1** | alpha/beta fold hydrolase | *Thiomonas delicata* |
| **WP_233468069.1** | alpha/beta fold hydrolase | *Hydrogenophilus thermoluteolus* |

| | | |
|---|---|---|
| **OZB45810.1** | MAG: poly-beta-hydroxybutyrate polymerase | *Thiomonas sp. 15-66-11* |
| **WP_265136348.1** | alpha/beta fold hydrolase | *Caldimonas thermodepolymerans* |
| **WP_104357307.1** | alpha/beta fold hydrolase | *Caldimonas thermodepolymerans* |
| **SBP89383.1** | putative Poly-beta-hydroxybutyrate polymerase | *Thiomonas delicata* |
| **WP_132766326.1** | alpha/beta fold hydrolase | *Caldimonas thermodepolymerans* |
| **WP_112488320.1** | alpha/beta fold hydrolase | *Thiomonas sp. X19* |
| **WP_018914728.1** | alpha/beta fold hydrolase | *Thiomonas sp. FB-6* |
| **WP_076167847.1** | hypothetical protein | *Amycolatopsis coloradensis* |
| **MBD0358487.1** | alpha/beta fold hydrolase | *Rubrobacter sp.* |
| **OBS30746.1** | hypothetical protein A9O67_07195 | *Tepidimonas fonticaldi* |
| **WP_143969726.1** | poly-beta-hydroxybutyrate polymerase N-terminal domain-containing protein | *Tepidimonas fonticaldi* |
| **WP_156682287.1** | poly-beta-hydroxybutyrate polymerase N-terminal domain-containing protein | *Tepidimonas fonticaldi* |
| **WP_143899688.1** | poly-beta-hydroxybutyrate polymerase N-terminal domain-containing protein | *Tepidimonas thermarum* |
| **WP_143892729.1** | poly-beta-hydroxybutyrate polymerase N-terminal domain-containing protein | *Tepidimonas sediminis* |
| **WP_244162576.1** | poly-beta-hydroxybutyrate polymerase | *Amycolatopsis regifaucium* |
| **KZB85987.1** | hypothetical protein AVL48_27685 | *Amycolatopsis regifaucium* |
| **TSE36391.1** | Poly(3-hydroxyalkanoate) polymerase subunit PhaC | *Tepidimonas charontis* |
| **WP_144327062.1** | poly-beta-hydroxybutyrate polymerase N-terminal domain-containing protein | *Tepidimonas charontis* |
| **MBN1811673.1** | alpha/beta fold hydrolase | *Anaerolineae bacterium* |
| **MCB1175590.1** | alpha/beta fold hydrolase | *Leptospiraceae bacterium* |
| **MBA4115524.1** | class III poly(R)-hydroxyalkanoic acid synthase subunit PhaC | *Rubrobacter sp.* |
| **WP_132691807.1** | class III poly(R)-hydroxyalkanoic acid synthase subunit PhaC | *Rubrobacter taiwanensis* |
| **MBX3014313.1** | class III poly(R)-hydroxyalkanoic acid synthase subunit PhaC | *Caldilineaceae bacterium* |
| **WP_166397552.1** | class III poly(R)-hydroxyalkanoic acid synthase subunit PhaC | *Rubrobacter marinus* |
| **MCB9117856.1** | class III poly(R)-hydroxyalkanoic acid synthase subunit PhaC | *Caldilineaceae bacterium* |
| **MCB0155473.1** | class III poly(R)-hydroxyalkanoic acid synthase subunit PhaC | *Anaerolineae bacterium* |
| **MBA2692587.1** | class III poly(R)-hydroxyalkanoic acid synthase subunit PhaC | *Rubrobacter sp.* |
| **MCB0039881.1** | class III poly(R)-hydroxyalkanoic acid synthase subunit PhaC | *Caldilinea sp.* |
| **WP_219974539.1** | class III poly(R)-hydroxyalkanoic acid synthase subunit PhaC | *Rubrobacter xylanophilus* |
| **WP_077732268.1** | MULTISPECIES: class III poly(R)-hydroxyalkanoic acid synthase subunit PhaC | *unclassified Methylocaldum* |
| **MBA3422947.1** | class III poly(R)-hydroxyalkanoic acid synthase subunit PhaC | *Rubrobacter sp.* |
| **WP_266096606.1** | alpha/beta fold hydrolase | *Rubrobacter marinus* |
| **MBA3388422.1** | class III poly(R)-hydroxyalkanoic acid synthase subunit PhaC | *Rubrobacter sp.* |
| **WP_011565326.1** | class III poly(R)-hydroxyalkanoic acid synthase subunit PhaC | *Rubrobacter xylanophilus* |
| **MCL4300363.1** | class III poly(R)-hydroxyalkanoic acid synthase subunit PhaC | *Anaerolineae bacterium* |
| **WP_036269290.1** | class III poly(R)-hydroxyalkanoic acid synthase subunit PhaC | *Methylocaldum szegediense* |
| **WP_119628989.1** | class III poly(R)-hydroxyalkanoic acid synthase subunit PhaC | *Methylocaldum marinum* |
| **MCE7983580.1** | class III poly(R)-hydroxyalkanoic acid synthase subunit PhaC | *Caldilinea sp. CFX5* |
| **MCB0083735.1** | class III poly(R)-hydroxyalkanoic acid synthase subunit PhaC | *Caldilineaceae bacterium* |
| **MCB0145186.1** | class III poly(R)-hydroxyalkanoic acid synthase subunit PhaC | *Caldilineaceae bacterium* |

| | | |
|---|---|---|
| **WP_143527769.1** | class III poly(R)-hydroxyalkanoic acid synthase subunit PhaC | *Rubrobacter xylanophilus* |
| **WP_047865909.1** | class III poly(R)-hydroxyalkanoic acid synthase subunit PhaC | *Rubrobacter aplysinae* |
| **MBW7884932.1** | class III poly(R)-hydroxyalkanoic acid synthase subunit PhaC | *Caldilineaceae bacterium* |
| **MBE2268011.1** | class III poly(R)-hydroxyalkanoic acid synthase subunit PhaC | *Anaerolinea sp.* |
| **OPX86172.1** | MAG: Poly-beta-hydroxybutyrate polymerase | *Pelotomaculum sp. PtaB.Bin104* |
| **WP_181538453.1** | class III poly(R)-hydroxyalkanoic acid synthase subunit PhaC | *Anoxybacillus calidus* |
| **OPX86175.1** | MAG: Poly-beta-hydroxybutyrate polymerase | *Pelotomaculum sp. PtaB.Bin104* |
| **WP_111643468.1** | class III poly(R)-hydroxyalkanoic acid synthase subunit PhaC | *Anoxybacillus vitaminiphilus* |
| **WP_070111171.1** | class III poly(R)-hydroxyalkanoic acid synthase subunit PhaC | *Clostridium acetireducens* |
| **MCC6798982.1** | alpha/beta fold hydrolase | *Anaerolineae bacterium* |
| **MCB0129538.1** | class III poly(R)-hydroxyalkanoic acid synthase subunit PhaC | *Caldilineaceae bacterium* |
| **WP_026964144.1** | class III poly(R)-hydroxyalkanoic acid synthase subunit PhaC | *Alicyclobacillus pomorum* |
| **WP_038682548.1** | class III poly(R)-hydroxyalkanoic acid synthase subunit PhaC | *Rubrobacter radiotolerans* |
| **WP_074783018.1** | class III poly(R)-hydroxyalkanoic acid synthase subunit PhaC | *Clostridium homopropionicum* |
| **MBD0358486.1** | class III poly(R)-hydroxyalkanoic acid synthase subunit PhaC | *Rubrobacter sp.* |
| **WP_166173905.1** | class III poly(R)-hydroxyalkanoic acid synthase subunit PhaC | *Rubrobacter tropicus* |
| **WP_166397557.1** | class III poly(R)-hydroxyalkanoic acid synthase subunit PhaC | *Rubrobacter marinus* |
| **MCB9161166.1** | class III poly(R)-hydroxyalkanoic acid synthase subunit PhaC | *Caldilineaceae bacterium* |
| **MCB9177621.1** | class III poly(R)-hydroxyalkanoic acid synthase subunit PhaC | *Caldilineae bacterium* |
| **MCB0158739.1** | class III poly(R)-hydroxyalkanoic acid synthase subunit PhaC | *Caldilineaceae bacterium* |
| **WP_052222780.1** | class III poly(R)-hydroxyalkanoic acid synthase subunit PhaC | *Clostridium homopropionicum* |
| **MCD4687217.1** | class III poly(R)-hydroxyalkanoic acid synthase subunit PhaC | *Anaerolineae bacterium* |
| **MCG3210572.1** | Poly(3-hydroxyalkanoate) polymerase subunit PhaC | *Anaerolineae bacterium* |
| **WP_119069243.1** | class III poly(R)-hydroxyalkanoic acid synthase subunit PhaC | *Rubrobacter indicoceani* |
| **MBD0358830.1** | class III poly(R)-hydroxyalkanoic acid synthase subunit PhaC | *Rubrobacter sp.* |
| **MBA4179263.1** | poly-beta-hydroxybutyrate polymerase | *Anaerolinea sp.* |
| **THB64347.1** | alpha/beta fold hydrolase | *Spirochaetaceae bacterium* |
| **MCD4687216.1** | class III poly(R)-hydroxyalkanoic acid synthase subunit PhaC | *Anaerolineae bacterium* |
| **MBE7468601.1** | class III poly(R)-hydroxyalkanoic acid synthase subunit PhaC | *Anaerolineales bacterium* |
| **NJN93336.1** | class III poly(R)-hydroxyalkanoic acid synthase subunit PhaC | *Anaerolineales bacterium* |
| **MBN8636024.1** | class III poly(R)-hydroxyalkanoic acid synthase subunit PhaC | *Anaerolineae bacterium* |
| **MCK6625714.1** | class III poly(R)-hydroxyalkanoic acid synthase subunit PhaC | *Anaerolineae bacterium* |
| **MBA2441667.1** | class III poly(R)-hydroxyalkanoic acid synthase subunit PhaC | *Rubrobacter sp.* |
| **MCB0209481.1** | class III poly(R)-hydroxyalkanoic acid synthase subunit PhaC | *Anaerolineae bacterium* |
| **MCB0170077.1** | class III poly(R)-hydroxyalkanoic acid synthase subunit PhaC | *Anaerolineae bacterium* |
| **MCL4238390.1** | alpha/beta fold hydrolase | *Anaerolineae bacterium* |

| | | |
|---|---|---|
| **MBA3474707.1** | class III poly(R)-hydroxyalkanoic acid synthase subunit PhaC | *Rubrobacter sp.* |
| **OHD10767.1** | hypothetical protein A2Y41_00920 | *Spirochaetes bacterium GWB1_36_13* |
| **WP_208649453.1** | class III poly(R)-hydroxyalkanoic acid synthase subunit PhaC | *Ureibacillus thermophilus* |
| **WP_273841781.1** | class III poly(R)-hydroxyalkanoic acid synthase subunit PhaC | *Rubrobacter calidifluminis* |
| **MBN2044693.1** | class III poly(R)-hydroxyalkanoic acid synthase subunit PhaC | *Anaerolineales bacterium* |
| **WP_119071681.1** | alpha/beta fold hydrolase | *Aggregatilinea lenta* |
| **MBP8974062.1** | alpha/beta fold hydrolase | *Anaerolineae bacterium* |
| **QRG70492.1** | class III poly(R)-hydroxyalkanoic acid synthase subunit PhaC | *Brevibacillus choshinensis* |
| **WP_238933592.1** | class III poly(R)-hydroxyalkanoic acid synthase subunit PhaC | *Brevibacillus choshinensis* |
| **MCB0193585.1** | class III poly(R)-hydroxyalkanoic acid synthase subunit PhaC | *Anaerolineae bacterium* |
| **WP_256644884.1** | class III poly(R)-hydroxyalkanoic acid synthase subunit PhaC | *Thermomonas paludicola* |
| **MBA4116379.1** | alpha/beta fold hydrolase | *Rubrobacter sp.* |
| **MBB5147750.1** | polyhydroxyalkanoate synthase | *Ureibacillus thermosphaericus* |
| **WP_273887813.1** | class III poly(R)-hydroxyalkanoic acid synthase subunit PhaC | *Rubrobacter naiadicus* |
| **WP_038683188.1** | alpha/beta fold hydrolase | *Rubrobacter radiotolerans* |
| **WP_210730726.1** | class III poly(R)-hydroxyalkanoic acid synthase subunit PhaC | *Ureibacillus thermosphaericus* |
| **WP_016836885.1** | class III poly(R)-hydroxyalkanoic acid synthase subunit PhaC | *Ureibacillus thermosphaericus* |
| **WP_096550753.1** | class III poly(R)-hydroxyalkanoic acid synthase subunit PhaC | *Ureibacillus thermosphaericus* |
| **WP_147666195.1** | class III poly(R)-hydroxyalkanoic acid synthase subunit PhaC | *Cerasibacillus terrae* |
| **WP_246045766.1** | class III poly(R)-hydroxyalkanoic acid synthase subunit PhaC | *Ureibacillus terrenus* |
| **TQE92436.1** | class III poly(R)-hydroxyalkanoic acid synthase subunit PhaC | *Ureibacillus terrenus* |
| **WP_173680423.1** | class III poly(R)-hydroxyalkanoic acid synthase subunit PhaC | *Clostridium tetanomorphum* |
| **KGM42515.1** | poly-beta-hydroxybutyrate polymerase | *Alkalispirochaeta odontotermitis* |
| **MBA3388109.1** | hypothetical protein | *Rubrobacter sp.* |
| **WP_173680355.1** | class III poly(R)-hydroxyalkanoic acid synthase subunit PhaC | *Clostridium tetanomorphum* |
| **WP_111947812.1** | class III poly(R)-hydroxyalkanoic acid synthase subunit PhaC | *Clostridium tetanomorphum* |
| **WP_242846772.1** | class III poly(R)-hydroxyalkanoic acid synthase subunit PhaC | *Clostridium homopropionicum* |
| **WP_122916151.1** | class III poly(R)-hydroxyalkanoic acid synthase subunit PhaC | *Brevibacillus fluminis* |
| **WP_035152738.1** | class III poly(R)-hydroxyalkanoic acid synthase subunit PhaC | *Clostridium tetanomorphum* |
| **WP_166177073.1** | alpha/beta fold hydrolase | *Rubrobacter tropicus* |
| **HBQ29258.1** | class III poly(R)-hydroxyalkanoic acid synthase subunit PhaC | *Desulfotomaculum sp.* |
| **HAG08735.1** | class III poly(R)-hydroxyalkanoic acid synthase subunit PhaC | *Desulfotomaculum sp.* |
| **WP_209795125.1** | class III poly(R)-hydroxyalkanoic acid synthase subunit PhaC | *Clostridium tetanomorphum* |
| **WP_238021877.1** | class III poly(R)-hydroxyalkanoic acid synthase subunit PhaC | *Clostridium cochlearium* |
| **WP_187570609.1** | class III poly(R)-hydroxyalkanoic acid synthase subunit PhaC | *Thermomonas brevis* |
| **WP_278428054.1** | class III poly(R)-hydroxyalkanoic acid synthase subunit PhaC | *Clostridium cochlearium* |
| **HAU30729.1** | class III poly(R)-hydroxyalkanoic acid synthase subunit PhaC | *Desulfotomaculum sp.* |
| **KUK52556.1** | MAG: Alpha/beta hydrolase fold protein | *Desulfotomaculum sp. 46_296* |
| **WP_153975441.1** | class III poly(R)-hydroxyalkanoic acid synthase subunit PhaC | *Thermochromatium tepidum* |
| **MBO8157996.1** | alpha/beta fold hydrolase | *Thermosyntropha sp.* |

| | | |
|---|---|---|
| **WP_240127068.1** | class III poly(R)-hydroxyalkanoic acid synthase subunit PhaC | *Thermomonas alba* |
| **WP_223629487.1** | class III poly(R)-hydroxyalkanoic acid synthase subunit PhaC | *Thermomonas beijingensis* |
| **MBE7551318.1** | class III poly(R)-hydroxyalkanoic acid synthase subunit PhaC | *Anaerolineales bacterium* |
| **MCA9940934.1** | alpha/beta fold hydrolase | *Anaerolineales bacterium* |
| **WP_190281238.1** | class III poly(R)-hydroxyalkanoic acid synthase subunit PhaC | *Thermomonas sp. XSG* |
| **WP_027624860.1** | class III poly(R)-hydroxyalkanoic acid synthase subunit PhaC | *Clostridium lundense* |
| **MCB9435617.1** | class III poly(R)-hydroxyalkanoic acid synthase subunit PhaC | *Anaerolineales bacterium* |
| **PMP81362.1** | class III poly(R)-hydroxyalkanoic acid synthase subunit PhaC | *Roseiflexus castenholzii* |
| **WP_273388233.1** | class III poly(R)-hydroxyalkanoic acid synthase subunit PhaC | *Thermomonas hydrothermalis* |
| **PKL15222.1** | hypothetical protein CVV50_00790 | *Spirochaetae bacterium HGW-Spirochaetae-6* |
| **MBO9341583.1** | alpha/beta fold hydrolase | *Roseiflexus sp.* |
| **WP_011959221.1** | alpha/beta fold hydrolase | *Roseiflexus sp. RS-1* |
| **WP_072755904.1** | class III poly(R)-hydroxyalkanoic acid synthase subunit PhaC | *Thermomonas hydrothermalis* |
| **MBV2209965.1** | class III poly(R)-hydroxyalkanoic acid synthase subunit PhaC | *Thermomonas sp.* |
| **WP_114958938.1** | class III poly(R)-hydroxyalkanoic acid synthase subunit PhaC | *Thermomonas haemolytica* |
| **WP_011997646.1** | alpha/beta fold hydrolase | *Roseiflexus castenholzii* |
| **MCS6840654.1** | alpha/beta fold hydrolase | *Roseiflexus sp.* |
| **WP_028840045.1** | class III poly(R)-hydroxyalkanoic acid synthase subunit PhaC | *Thermomonas fusca* |
| **WP_138348483.1** | class III poly(R)-hydroxyalkanoic acid synthase subunit PhaC | *Thermomonas fusca* |
| **MCS6938942.1** | alpha/beta fold hydrolase | *Roseiflexus sp.* |
| **MBO9325172.1** | alpha/beta fold hydrolase | *Roseiflexus sp.* |
| **KAJ52316.1** | poly(R)-hydroxyalkanoic acid synthase subunit PhaC | *Clostridium tetanomorphum DSM 665* |
| **WP_036183957.1** | class III poly(R)-hydroxyalkanoic acid synthase subunit PhaC | *Ureibacillus manganicus* |
| **OHD07979.1** | hypothetical protein A2Y41_13805 | *Spirochaetes bacterium GWB1_36_13* |
| **OYV39239.1** | MAG: poly-beta-hydroxybutyrate polymerase, partial | *Thiomonas sp. 20-64-5* |
| **MBK6417342.1** | class III poly(R)-hydroxyalkanoic acid synthase subunit PhaC | *Thermomonas sp.* |
| **MBP7787837.1** | class III poly(R)-hydroxyalkanoic acid synthase subunit PhaC | *Thermomonas sp.* |
| **MBK6332836.1** | class III poly(R)-hydroxyalkanoic acid synthase subunit PhaC | *Thermomonas sp.* |
| **MCR6496130.1** | class III poly(R)-hydroxyalkanoic acid synthase subunit PhaC | *Thermomonas sp. S9* |
| **MBP9696220.1** | class III poly(R)-hydroxyalkanoic acid synthase subunit PhaC | *Thermomonas sp.* |
| **WP_073089411.1** | alpha/beta fold hydrolase | *Thermosyntropha lipolytica* |
| **WP_259304761.1** | class III poly(R)-hydroxyalkanoic acid synthase subunit PhaC | *Thermomonas sp. S9* |
| **WP_066789608.1** | alpha/beta fold hydrolase | *Chloroflexus islandicus* |
| **MBK9668690.1** | class III poly(R)-hydroxyalkanoic acid synthase subunit PhaC | *Thermomonas sp.* |
| **WP_187552196.1** | class III poly(R)-hydroxyalkanoic acid synthase subunit PhaC | *Thermomonas carbonis* |
| **OHD06989.1** | hypothetical protein A2Y41_11120 | *Spirochaetes bacterium GWB1_36_13* |
| **MBK6924577.1** | class III poly(R)-hydroxyalkanoic acid synthase subunit PhaC | *Thermomonas sp.* |
| **WP_012615932.1** | alpha/beta fold hydrolase | *Chloroflexus aggregans* |
| **PMP76775.1** | class III poly(R)-hydroxyalkanoic acid synthase subunit PhaC | *Chloroflexus aggregans* |
| **TXH64290.1** | MAG: class III poly(R)-hydroxyalkanoic acid synthase subunit PhaC | *Thermomonas sp.* |

| | | |
|---|---|---|
| **WP_240099333.1** | class III poly(R)-hydroxyalkanoic acid synthase subunit PhaC | *Thermomonas flagellata* |
| **WP_012259104.1** | alpha/beta fold hydrolase | *Chloroflexus aurantiacus* |
| **HCL55721.1** | class III poly(R)-hydroxyalkanoic acid synthase subunit PhaC | *Spirochaetia bacterium* |
| **WP_271678800.1** | class III poly(R)-hydroxyalkanoic acid synthase subunit PhaC | *Thermomonas sp. 2C3345* |
| **WP_031458694.1** | alpha/beta fold hydrolase | *Chloroflexus sp. MS-G* |
| **WP_028456980.1** | alpha/beta fold hydrolase | *Chloroflexus sp. Y-396-1* |
| **GIV94657.1** | MAG: poly(R)-hydroxyalkanoic acid synthase | *Chloroflexus sp.* |
| **MBO9339101.1** | alpha/beta fold hydrolase | *Chloroflexus sp.* |
| **MCO5054012.1** | class III poly(R)-hydroxyalkanoic acid synthase subunit PhaC | *Thermomonas sp.* |
| **MBQ4330322.1** | class III poly(R)-hydroxyalkanoic acid synthase subunit PhaC | *Spirochaetaceae bacterium* |
| **MCA9940935.1** | alpha/beta fold hydrolase | *Anaerolineales bacterium* |
| **MBO5125060.1** | class III poly(R)-hydroxyalkanoic acid synthase subunit PhaC | *Spirochaetaceae bacterium* |
| **MBO5690322.1** | class III poly(R)-hydroxyalkanoic acid synthase subunit PhaC | *Spirochaetaceae bacterium* |
| **MBP3416397.1** | class III poly(R)-hydroxyalkanoic acid synthase subunit PhaC | *Spirochaetaceae bacterium* |
| **MBO5730312.1** | class III poly(R)-hydroxyalkanoic acid synthase subunit PhaC | *Treponema sp.* |
| **WP_166056882.1** | class III poly(R)-hydroxyalkanoic acid synthase subunit PhaC | *Thermomonas sp. HDW16* |
| **MBZ0086782.1** | class III poly(R)-hydroxyalkanoic acid synthase subunit PhaC | *Thermomonas sp.* |
| **WP_139715061.1** | class III poly(R)-hydroxyalkanoic acid synthase subunit PhaC | *Thermomonas aquatica* |
| **MBN1811671.1** | class III poly(R)-hydroxyalkanoic acid synthase subunit PhaC | *Anaerolineae bacterium* |
| **MCC7097066.1** | class III poly(R)-hydroxyalkanoic acid synthase subunit PhaC | *Thermomonas sp.* |
| **MBL6982748.1** | alpha/beta fold hydrolase | *Anaerolineales bacterium* |
| **MCB1175800.1** | alpha/beta fold hydrolase | *Leptospiraceae bacterium* |
| **QOS92875.1** | alpha/beta fold hydrolase | *Brevibacillus sp. JNUCC-41* |
| **MCB9135562.1** | alpha/beta fold hydrolase | *Anaerolineales bacterium* |
| **NUM46194.1** | alpha/beta fold hydrolase | *Anaerolineales bacterium* |
| **MCA9991047.1** | alpha/beta fold hydrolase | *Anaerolineales bacterium* |
| **MCB9138516.1** | class III poly(R)-hydroxyalkanoic acid synthase subunit PhaC | *Caldilineaceae bacterium* |
| **MCB0193330.1** | class III poly(R)-hydroxyalkanoic acid synthase subunit PhaC | *Anaerolineae bacterium* |
| **WP_052222537.1** | alpha/beta fold hydrolase | *Clostridium homopropionicum* |
| **MCA9934284.1** | alpha/beta fold hydrolase | *Anaerolineales bacterium* |
| **MCA9926268.1** | alpha/beta fold hydrolase | *Anaerolineales bacterium* |
| **MCA9927351.1** | alpha/beta fold hydrolase | *Anaerolineales bacterium* |
| **MCB9099623.1** | alpha/beta fold hydrolase | *Anaerolineales bacterium* |
| **MCB0178194.1** | alpha/beta fold hydrolase | *Anaerolineae bacterium* |
| **WP_073090307.1** | alpha/beta fold hydrolase | *Thermosyntropha lipolytica* |
| **MBA3953302.1** | alpha/beta fold hydrolase | *Rubrobacter sp.* |
| **MBO8158156.1** | alpha/beta fold hydrolase | *Thermosyntropha sp.* |
| **MCB0104875.1** | class III poly(R)-hydroxyalkanoic acid synthase subunit PhaC | *Caldilineaceae bacterium* |
| **MCB0093581.1** | class III poly(R)-hydroxyalkanoic acid synthase subunit PhaC | *Caldilineaceae bacterium* |
| **KUK53159.1** | MAG: hypothetical protein XD78_1440 | *Desulfotomaculum sp. 46_296* |
| **GHG72360.1** | alpha/beta hydrolase | *Amycolatopsis acidiphila* |
| **WP_144638073.1** | alpha/beta fold hydrolase | *Amycolatopsis acidiphila* |
| **WP_094862267.1** | alpha/beta fold hydrolase | *Amycolatopsis antarctica* |
| **HHV16637.1** | alpha/beta fold hydrolase | *Gelria sp.* |
| **WP_159720347.1** | MULTISPECIES: alpha/beta fold hydrolase | *Anoxybacillus* |
| **WP_080862764.1** | alpha/beta fold hydrolase | *Anoxybacillus sp. UARK-01* |
| **WP_066148787.1** | MULTISPECIES: alpha/beta fold hydrolase | *Anoxybacillus* |

| | | |
|---|---|---|
| **WP_212388024.1** | alpha/beta fold hydrolase | *Anoxybacillus rupiensis* |
| **WP_236789170.1** | alpha/beta fold hydrolase | *Amycolatopsis sp. GM8* |
| **MCB0057522.1** | alpha/beta fold hydrolase | *Caldilineaceae bacterium* |
| **WP_209662253.1** | alpha/beta hydrolase | *Amycolatopsis magusensis* |
| **WP_052660947.1** | MULTISPECIES: alpha/beta fold hydrolase | *unclassified Anoxybacillus* |
| **HDP79532.1** | alpha/beta hydrolase | *Spirochaetota bacterium* |
| **WP_094161467.1** | alpha/beta fold hydrolase | *Thiomonas delicata* |

**Supplementary Table S3**: Filtered list of putative PhaC sequences in archaeal moderate thermophiles. This list is filtered on the lipase-like box location and the conservation of the hyperconserved residues.

| Accession | Annotation | Microorganism |
|---|---|---|
| **WP_006167436.1** | class III poly(R)-hydroxyalkanoic acid synthase subunit PhaC | *Natrialba chahannaoensis* |
| **WP_228717180.1** | MULTISPECIES: class III poly(R)-hydroxyalkanoic acid synthase subunit PhaC | *Haloferax* |
| **KAB1188150.1** | class III poly(R)-hydroxyalkanoic acid synthase subunit PhaC | *Haloferax sp. CBA1149* |
| **WP_008321928.1** | class III poly(R)-hydroxyalkanoic acid synthase subunit PhaC | *Haloferax elongans* |
| **WP_074796499.1** | class III poly(R)-hydroxyalkanoic acid synthase subunit PhaC | *Haloferax larsenii* |
| **WP_151110288.1** | MULTISPECIES: class III poly(R)-hydroxyalkanoic acid synthase subunit PhaC | *Haloferax* |
| **WP_249361452.1** | class III poly(R)-hydroxyalkanoic acid synthase subunit PhaC | *Haloterrigena sp. H1* |
| **WP_007542254.1** | class III poly(R)-hydroxyalkanoic acid synthase subunit PhaC | *Haloferax larsenii* |
| **TMT87062.1** | class III poly(R)-hydroxyalkanoic acid synthase subunit PhaC | *Haloterrigena sp. H1* |
| **TMT81407.1** | class III poly(R)-hydroxyalkanoic acid synthase subunit PhaC | *Haloterrigena sp. H1* |
| **WP_258303609.1** | class III poly(R)-hydroxyalkanoic acid synthase subunit PhaC | *Haloferax larsenii* |
| **WP_004060696.1** | class III poly(R)-hydroxyalkanoic acid synthase subunit PhaC | *Haloferax mediterranei* |
| **WP_121597719.1** | class III poly(R)-hydroxyalkanoic acid synthase subunit PhaC | *Halorubrum sp. Atlit-26R* |
| **WP_207289079.1** | class III poly(R)-hydroxyalkanoic acid synthase subunit PhaC | *Haloterrigena alkaliphila* |
| **WP_149081248.1** | class III poly(R)-hydroxyalkanoic acid synthase subunit PhaC | *Natrialba swarupiae* |
| **WP_004056138.1** | class III poly(R)-hydroxyalkanoic acid synthase subunit PhaC | *Haloferax mediterranei* |
| **WP_008320237.1** | class III poly(R)-hydroxyalkanoic acid synthase subunit PhaC | *Haloferax mucosum* |
| **WP_160047113.1** | class III poly(R)-hydroxyalkanoic acid synthase subunit PhaC | *Natrialba sp. INN-245* |
| **WP_008894642.1** | class III poly(R)-hydroxyalkanoic acid synthase subunit PhaC | *Haloterrigena salina* |
| **WP_004060157.1** | class III poly(R)-hydroxyalkanoic acid synthase subunit PhaC | *Haloferax mediterranei* |
| **WP_174679405.1** | class III poly(R)-hydroxyalkanoic acid synthase subunit PhaC | *Haloterrigena gelatinilytica* |
| **WP_012943615.1** | class III poly(R)-hydroxyalkanoic acid synthase subunit PhaC | *Haloterrigena turkmenica* |
| **WP_126661474.1** | class III poly(R)-hydroxyalkanoic acid synthase subunit PhaC | *Haloterrigena salifodinae* |
| **WP_204747298.1** | class III poly(R)-hydroxyalkanoic acid synthase subunit PhaC | *Haloterrigena salifodinae* |
| **RDZ56238.1** | class III poly(R)-hydroxyalkanoic acid synthase subunit PhaC | *Haloferax sp. Atlit-10N* |
| **WP_233547734.1** | class III poly(R)-hydroxyalkanoic acid synthase subunit PhaC | *Haloferax sp. Atlit-10N* |

| | | |
|---|---|---|
| **WP_137683212.1** | poly(3-hydroxyalkanoate) polymerase subunit PhaC | *Haloarcula mannanilytica* |
| **WEL22451.1** | poly[(R)-3-hydroxyalkanoate] polymerase subunit PhaC | *Halorhabdus sp. BNX81* |
| **WP_275882327.1** | class III poly(R)-hydroxyalkanoic acid synthase subunit PhaC | *Halorhabdus sp. BNX81* |
| **WP_136689751.1** | class III poly(R)-hydroxyalkanoic acid synthase subunit PhaC | *Halorhabdus amylolytica* |
| **WP_165897347.1** | poly(3-hydroxyalkanoate) polymerase subunit PhaC | *Haloarcula sp. JP-Z28* |
| **WP_127013871.1** | MULTISPECIES: poly(3-hydroxyalkanoate) polymerase subunit PhaC | *Haloarcula* |
| **WP_004960314.1** | poly(3-hydroxyalkanoate) polymerase subunit PhaC | *Haloarcula sinaiiensis* |
| **WEL18563.1** | poly[(R)-3-hydroxyalkanoate] polymerase subunit PhaC | *Halorhabdus sp. SVX81* |
| **WP_275737459.1** | class III poly(R)-hydroxyalkanoic acid synthase subunit PhaC | *Halorhabdus sp. SVX81* |
| **WP_101348954.1** | MULTISPECIES: poly(3-hydroxyalkanoate) polymerase subunit PhaC | *Haloarcula* |
| **WP_050037370.1** | poly(3-hydroxyalkanoate) polymerase subunit PhaC | *Haloarcula sp. CBA1115* |
| **QPF21196.1** | poly(3-hydroxyalkanoate) synthase subunit C, partial | *Haloarcula sp.* |
| **QIO21410.1** | class III poly(R)-hydroxyalkanoic acid synthase subunit PhaC | *Haloarcula sp. JP-L23* |
| **WP_135665065.1** | class III poly(R)-hydroxyalkanoic acid synthase subunit PhaC | *Halorhabdus rudnickae* |
| **WP_151144736.1** | poly(3-hydroxyalkanoate) polymerase subunit PhaC | *Haloarcula sp. CBA1131* |
| **WP_058990864.1** | MULTISPECIES: poly(3-hydroxyalkanoate) polymerase subunit PhaC | *Haloarcula* |
| **WP_151102769.1** | poly(3-hydroxyalkanoate) polymerase subunit PhaC | *Haloarcula hispanica* |
| **WP_121512051.1** | poly(3-hydroxyalkanoate) polymerase subunit PhaC | *Haloarcula sp. Atlit-7R* |
| **WP_014039171.1** | poly(3-hydroxyalkanoate) polymerase subunit PhaC | *Haloarcula hispanica* |
| **WP_129755870.1** | poly(3-hydroxyalkanoate) polymerase subunit PhaC | *Haloarcula hispanica* |
| **WP_121302092.1** | poly(3-hydroxyalkanoate) polymerase subunit PhaC | *Haloarcula quadrata* |
| **WP_007187895.1** | poly(3-hydroxyalkanoate) polymerase subunit PhaC | *Haloarcula californiae* |
| **WP_008523978.1** | class III poly(R)-hydroxyalkanoic acid synthase subunit PhaC | *Halorhabdus tiamatea* |
| **WP_058993304.1** | poly(3-hydroxyalkanoate) polymerase subunit PhaC | *Haloarcula sp. CBA1127* |
| **WP_053966938.1** | poly(3-hydroxyalkanoate) polymerase subunit PhaC | *Haloarcula rubripromontorii* |
| **WP_011224492.1** | poly(3-hydroxyalkanoate) polymerase subunit PhaC | *Haloarcula marismortui* |
| **WP_008306972.1** | poly(3-hydroxyalkanoate) polymerase subunit PhaC | *Haloarcula amylolytica* |
| **WP_004516058.1** | poly(3-hydroxyalkanoate) polymerase subunit PhaC | *Haloarcula vallismortis* |
| **WP_151101215.1** | MULTISPECIES: poly(3-hydroxyalkanoate) polymerase subunit PhaC | *unclassified Haloarcula* |
| **WP_181687051.1** | class III poly(R)-hydroxyalkanoic acid synthase subunit PhaC | *Halorhabdus salina* |
| **WP_154552922.1** | MULTISPECIES: class III poly(R)-hydroxyalkanoic acid synthase subunit PhaC | *unclassified Halorhabdus* |
| **ABV71394.1** | poly(3-hydroxyalkanoate) synthase subunit PhaC | *Haloarcula hispanica* |

| | | |
|---|---|---|
| **WP_015789901.1** | class III poly(R)-hydroxyalkanoic acid synthase subunit PhaC | *Halorhabdus utahensis* |
| **WP_004590065.1** | poly(3-hydroxyalkanoate) polymerase subunit PhaC | *Haloarcula japonica* |
| **WP_170096459.1** | poly(3-hydroxyalkanoate) polymerase subunit PhaC | *Haloarcula argentinensis* |
| **WP_188975977.1** | poly(3-hydroxyalkanoate) polymerase subunit PhaC | *Haloarcula sebkhae* |
| **WP_188852231.1** | poly(3-hydroxyalkanoate) polymerase subunit PhaC | *Haloarcula salaria* |
| **EMA21044.1** | poly(3-hydroxyalkanoate) synthase | *Haloarcula argentinensis DSM 12282* |
| **WP_004061055.1** | alpha/beta fold hydrolase | *Haloferax mediterranei* |

**Supplementary Table S4**: Filtered list of putative PhaC sequences in extreme archaeal thermophiles. This list is filtered on the lipase-like box location and the conservation of the hyperconserved residues.

| Accession | Annotation | Microorganism |
|---|---|---|
| **WP_012965912.1** | alpha/beta fold hydrolase | *Ferroglobus placidus* |
| **MCC6027226.1** | alpha/beta fold hydrolase | *Archaeoglobus sp.* |
| **WP_169745351.1** | alpha/beta fold hydrolase | *Geoglobus ahangari* |
| **MCC6027983.1** | alpha/beta fold hydrolase | *Archaeoglobus sp.* |
| **WP_012965913.1** | alpha/beta fold hydrolase | *Ferroglobus placidus* |
| **WP_202319426.1** | alpha/beta fold hydrolase | *Archaeoglobus neptunius* |
| **HDD36547.1** | alpha/beta fold hydrolase | *Archaeoglobus veneficus* |

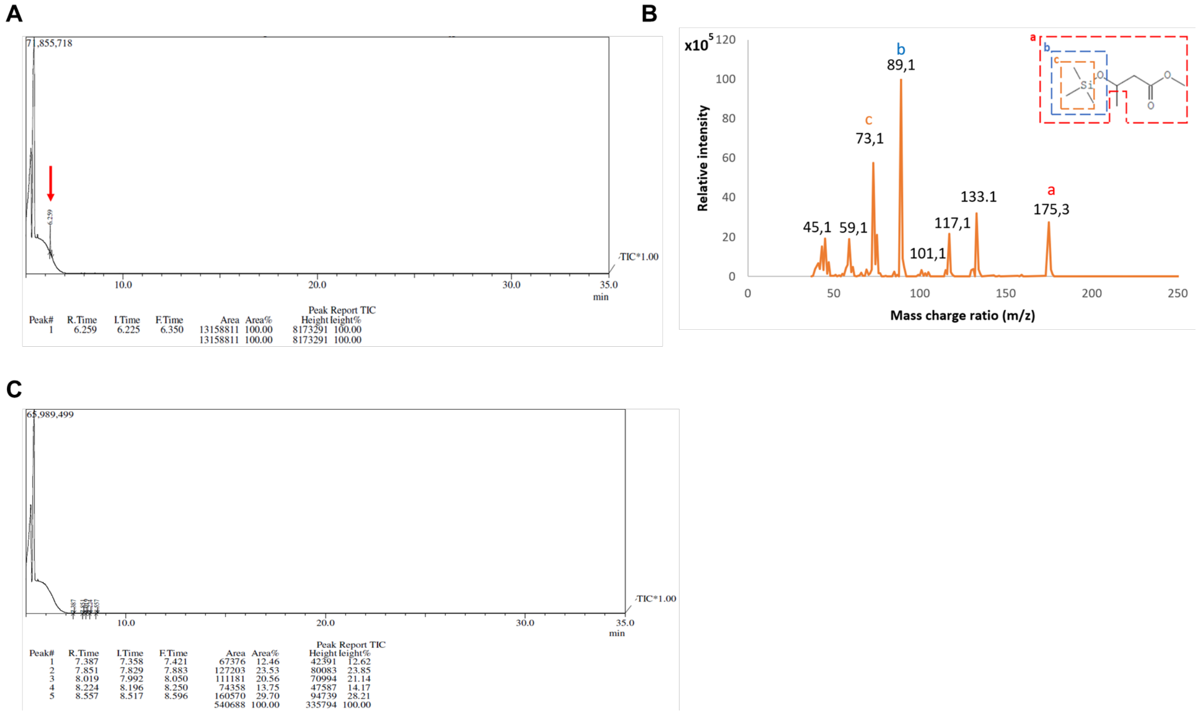


**Supplementary Figure S3**: GC-MS analysis. **(A)** GC chromatogram obtained from analysis of the polymer obtained from *Paracoccus denitrificans*. The x-axis represents the retention times, and the y-axis is a quantitative representation of the number of molecules that have a similar retention time. The red arrow represents the most important peak with a retention time characteristic of PHB. **(B)** MS spectrum for the extracted PHA from *P. denitrificans* that was obtained upon analyzing the main peak of the chromatogram with a retention time of 6.259 minutes. The x-axis represents the mass of the ions divided by their charge. The y-axis represents the signal intensity of the m/z peak. In the top right corner, the most important ions resulting from the electron impact and leading to this specific fingerprint are given. **(C)** GC chromatogram obtained from analysis of the sample of *T. thermophilus*. The characteristic peak of PHA was not found in this sample.

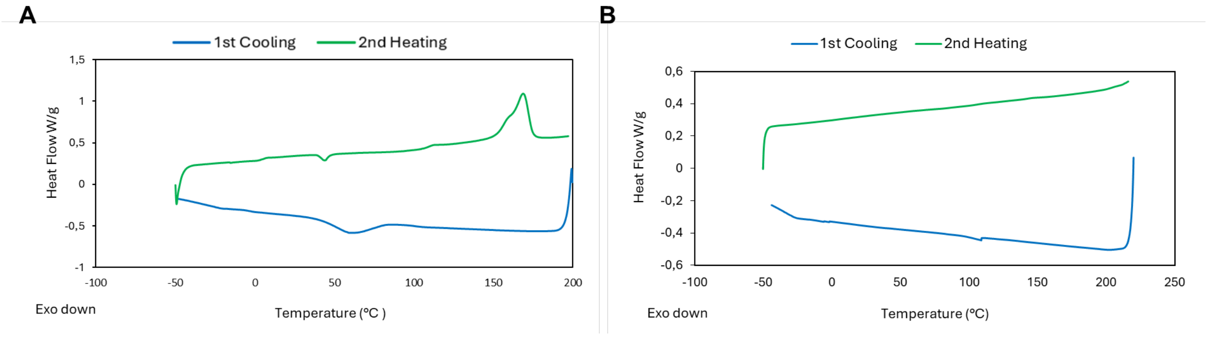


**Supplementary Figure S4:** DSC thermograms of the samples extracted from **(A)** *P. denitrificans* grown on MSM, showing the PHA characteristic thermal transitions with melting temperature ($T_m$) of 169°C and glass transition temperature ($T_g$) of 5°C. **(B)** *T. thermophilus* HB8, showing no thermal transitions demonstrating the absence of PHA in the extracted sample.

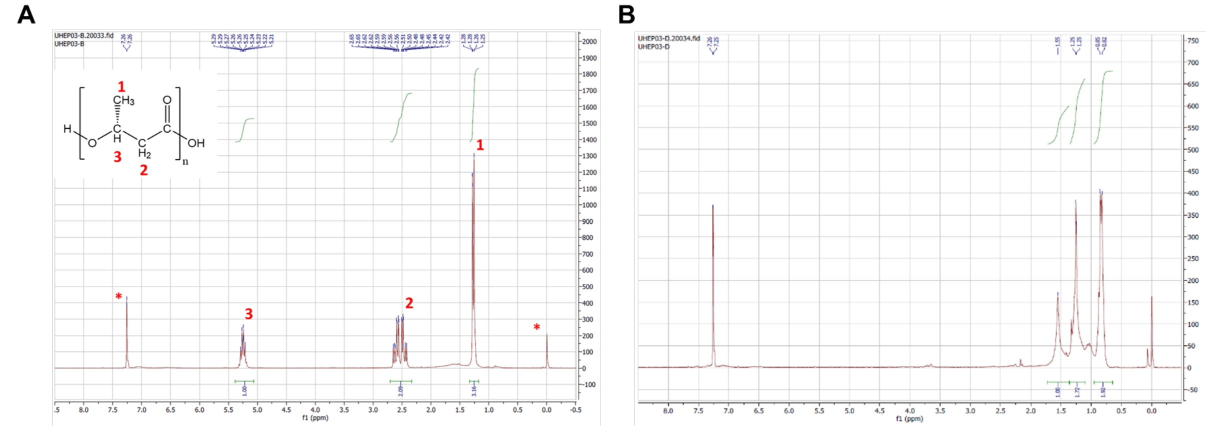


**Supplementary Figure S5**: [1]H NMR analysis performed for **(A)** a *P. denitrificans* sample and **(B)** a *T. thermophilus* sample. The Y-axis represents the signal intensity, and the X-axis represents the chemical shift of the protons in the sample. In the left upper corner of **(A)**, the structure of poly(3-hydroxybutyrate) is given with the protons assigned to the different peaks. Peaks that are labelled with an asterisk represent compounds unrelated to the polymer.
